# Cyclin B1-Cdk1 binding to MAD1 links nuclear pore disassembly to chromosomal stability

**DOI:** 10.1101/701474

**Authors:** Mark Jackman, Chiara Marcozzi, Mercedes Pardo, Lu Yu, Adam L. Tyson, Jyoti S. Choudhary, Jonathon Pines

**Affiliations:** The Institute of Cancer Research, 237 Fulham Road, London SW3 6JB

**Keywords:** Mitosis, Cyclin, Kinetochore, Spindle Assembly Checkpoint, TPR, MPS1

## Abstract

How the cell completely reorganises its architecture when it divides is a problem that has fascinated researchers for almost 150 years. We now know that the core regulatory machinery is highly conserved in eukaryotes but how these multiple protein kinases, protein phosphatases, and ubiquitin ligases are coordinated to remodel the cell in a matter of minutes remains a major question. Cyclin B-CDK is the primary kinase that drives mitotic remodelling and here we show that it is targeted to the nuclear pore complex (NPC) by binding an acidic face of the spindle assembly checkpoint protein, MAD1. This localised Cyclin B1-CDK1 activity coordinates NPC disassembly with kinetochore assembly: it is needed for the proper release of MAD1 from the embrace of TPR at the nuclear pore, which enables MAD1 to be recruited to kinetochores before nuclear envelope breakdown, thereby strengthening the spindle assembly checkpoint to maintain genomic stability.

## Introduction

The rapid and complete reorganisation of a cell at mitosis is one of the most striking events in cell biology, but we are only just beginning to understand how it is achieved. To understand the remarkable coordination required to remodel the interphase cell into a mitotic cell that is specialised to separate the genome equally into two daughter cells, we must elucidate the mechanisms by which the mitotic regulators disassemble interphase structures and promote the assembly of the mitotic apparatus. The conservation of much of the machinery through evolution has allowed us to identify that coordinated efforts of multiple protein kinases and phosphatases is required to remodel the cell. Chief amongst these are the activation of Cyclin B-CDK1 – the major mitotic kinase in almost all organisms studied to date – and the concomitant inhibition of its antagonistic PP2A-B55δ phosphatase (Castilho et al., 2009; Gharbi-Ayachi et al., 2010; Mochida et al., 2010). Together these drive the cell to enter mitosis. As the level of Cyclin B-CDK1 activity rises in the cell it triggers different events at different times (Gavet and Pines, 2010). But how this is achieved, and how the disassembly of interphase structures contributes to the assembly of mitosis-specific structures, are still largely unknown.

Although Cyclin B-CDK1 was identified as the major mitotic kinase in the 1980s (Arion et al., 1988; Dorée and Hunt, 2002; Dunphy and Newport, 1989; Labbe et al., 1988; Meijer et al., 1989; Minshull et al., 1989), and a plethora of crucial substrates identified since then (reviewed in Nigg, 1995; Wieser and Pines, 2015), it is remarkable that we still do not understand how it recognises its substrates. Our knowledge is limited to the minimal consensus sequence recognised by CDK1 (S/T-P, optimally in the context of basic residues) (Alexander et al., 2011; Bondt et al., 1993; Brown et al., 1999; Jeffrey et al., 1995), and evidence that its associated Cks subunit – which also binds to CDK2 – preferentially recognises phospho-threonines in a (F/I/L/P/V/W/Y-X-pT-P) consensus (McGrath et al., 2013). By contrast we know that the major interphase Cyclin-CDK complexes – Cyclins A and E, recognise many substrates through the Cy motif (RxL), which binds to the ‘hydrophobic patch’ on the first cyclin fold (Brown et al., 1999, 2007; Schulman et al., 1998), and that the D-type Cyclins have a LxCxE motif that recognises the Retinoblastoma protein (pRb) (Dowdy et al., 1993)

Elucidating how Cyclin B-CDK1 activity is directed to the right substrate at the right time is essential to understand how cells are remodelled because Cyclin B-CDK is both the essential trigger and the ‘workhorse’ of mitosis. Evidence for its role as the trigger of mitosis in mammals is that mouse embryos with a genetic deletion of Cyclin B1 (Brandeis et al., 1998) stop dividing around the 4-cell stage as soon as the maternal stock of Cyclin B1 is degraded (Strauss et al., 2018); these cells arrest in G2 phase and are unable to initiate mitosis (Strauss et al., 2018). To ensure that cells remain in mitosis Cyclin B-CDK1 phosphorylates and activates the Greatwall protein kinase, which generates an inhibitor of the PP2A-B55 phosphatase that antagonises Cyclin B-CDK1 in interphase (Castilho et al., 2009; Gharbi-Ayachi et al., 2010; Mochida et al., 2010). In its role as the workhorse of mitosis (Nigg, 1995), Cyclin B-CDK1 phosphorylates structural components throughout the cell including the nuclear lamins (Heald and McKeon, 1990; Peter et al., 1990), nuclear pore components (NPC) (Linder et al., 2017), condensins (Hirano, 2012) and cytoskeletal regulators such as Rho GEF ECT2 (Tatsumoto et al., 1999). Microtubule motors, endoplasmic reticulum and Golgi apparatus components are also extensively phosphorylated by CyclinB-CDK1 (Champion et al., 2017; Wieser and Pines, 2015). A crucial role for Cyclin B-CDK1 is to activate the Anaphase Promoting Complex/Cyclosome (APC/C): the ubiquitin ligase that will subsequently degrade Cyclin B itself (Fujimitsu et al., 2016; Golan et al., 2002; Lu et al., 2014; Passmore et al., 2005; Qiao et al., 2016) But Cyclin B-CDK is also required for the Spindle Assembly Checkpoint (SAC, (D’Angiolella et al., 2003; Hayward et al., 2019; Morin et al., 2012; Vázquez-Novelle et al., 2014) that keeps the APC/C from degrading Cyclin B (and the separase inhibitor, securin) until all the kinetochores are attached to the mitotic spindle and is essential for genomic stability (reviewed in Lara-Gonzalez et al., 2012; Musacchio and Salmon, 2007).

An insight into how one kinase can coordinate so many different events is that Cyclin B-CDK1 is targeted to different structures as the cell enters mitosis. Cyclin B-CDK is activated on centrosomes (Jackman et al., 2003) and a large fraction immediately moves into the nucleus over approximately 20 minutes preceding nuclear envelope breakdown (NEBD) (Gavet and Pines, 2010; Hagting et al., 1999; Pines and Hunter, 1991). Subsequently, Cyclin B1-CDK binds to the microtubules around the spindle caps, to chromosomes in early mitosis, and to unattached kinetochores (Hagting et al., 1999; Pines and Hunter, 1991; Bentley et al., 2007; Chen et al., 2008). These observations indicate that the localisation of Cyclin B-CDK1 may be an important determinant of how specific substrates are recognised at specific times.

The remodelling of the cell at mitosis raises another important question: when interphase macromolecular machines are disassembled in mitosis, do their components, or sub-complexes of their components, contribute to the function of newly assembled mitotic machines? For example, there is an intriguing connection between the NPC and kinetochores: when the NPC is disassembled at the end of prophase, several NPC components relocalise to kinetochores in mitosis, including the Nup 107-160 complex (Loïodice et al., 2004; Zuccolo et al., 2007), Nup358/RanBP2, and Crm1 (Dasso, 2006; Joseph et al., 2004) (reviewed in Forbes et al., 2015). Moreover, the MAD1 and MAD2 Spindle Assembly Checkpoint (SAC) proteins are prominently bound to the NPC in interphase (Chen et al., 1998; Lee et al., 2008); in mitosis these bind to unattached kinetochores to generate the Mitotic Checkpoint Complex (MCC, composed of MAD2, BUBR1, BUB3 and CDC20) that inhibits the APC/C to prevent premature sister chromatid separation and aneuploidy (London and Biggins, 2014; Moyle et al., 2014; Sudakin et al., 2001).

In budding yeast, which maintain a nuclear envelope during mitosis, MAD1 modulates nuclear transport in response to kinetochore detachment from microtubules (Cairo et al., 2013), but it is unclear whether MAD1 and MAD2 have an interphase role at the NPC in cells that breakdown their nuclear envelope in mitosis. Rodriguez-Bravo et al have proposed that in interphase the MAD1/MAD2 heterodimer at the nuclear pore in human cells can catalyse the production of the MCC (Rodriguez-Bravo et al., 2014) - before kinetochores are assembled from late G2 into mitosis (Gascoigne and Cheeseman, 2013) - and that this is required both to inhibit interphase APC/C and to generate sufficient MCC to inhibit the APC/C when it is fully activated at NEBD (Rodriguez-Bravo et al., 2014). The importance of this pathway is uncertain, however, since there are several other mechanisms that keep the APC/C in check in interphase: the Emi1 inhibitor and Cyclin A-CDK complexes both inhibit the Cdh1 coactivator (Di Fiore and Pines, 2007; Frye et al., 2013; Reimann et al., 2001; Sørensen et al., 2001), and phosphorylation by Cyclin A-CDK complexes prevents the CDC20 co-activator from binding the APC/C (Hein and Nilsson, 2016; Labit et al., 2012). Moreover, CDC20 cannot bind to the APC/C until an autoinhibitory loop of the APC1 subunit is phosphorylated by Cyclin-CDK and Plk1 kinases at mitosis (Qiao et al., 2016; Zhang et al., 2016). A recent report has proposed a further ‘timer’ mechanism that is activated after NEBD, whereby phosphorylation of the BUB1 protein by CDK1 and the MPS1 kinase recruits MAD1 to kinetochores to generate the MCC independently of the pathway that responds to microtubule attachment (Qian et al., 2017) Thus, the means by which the cell ensures that newly activated mitotic APC/C is kept inhibited until all kinetochores attach to microtubules is a matter of some debate.

Here we uncover a connection between disassembly of the NPC and the generation of a kinetochore competent for SAC-signalling that depends on the targeting of Cyclin B1 to MAD1. We show that the MAD1 protein binds to Cyclin B1 through an acidic patch in a predicted helical domain of MAD1, and that binding is required to recruit Cyclin B1 to unattached kinetochores. We further show that Cyclin B1 binding to MAD1 is important for the proper release of MAD1 from the nuclear pore and its timely recruitment to kinetochores before NEBD, and thus for the ability of kinetochores to generate a robust SAC signal in early mitosis and maintain genomic stability. Our findings furnish evidence for the importance of localised Cyclin B1-CDK1 activity in the coordinated reorganisation of the cell as it enters mitosis. We also provide a mechanism for how the cell coordinates activation of the APC/C with generating the MCC to keep the APC/C in check in early mitosis.

## Results

### Cyclin B1 binds to MAD1 through the acidic face of a helix encoded by exon 4

We sought to understand what controls the highly dynamic behaviour of Cyclin B1-CDK1 complexes as the cell enters mitosis; in particular, how Cyclin B1 is recruited to specific places in the cell at specific times. To identify binding partners, we immunoprecipitated Cyclin B1 from both normal diploid Retinal Pigment Epithelial (RPE) cells and from transformed HeLa cells, and analysed the co-precipitating proteins by mass spectrometry. In immunoprecipitates from both cell lines we found MAD1 as one of the most prominent proteins. We confirmed MAD1 as a major Cyclin B1-binding partner by immunoblotting. It co-immunoprecipitated with Cyclin B1 from cells in both G2 phase and mitosis (Figure 1A, see also Figures S1A, S1B), but we noticed that there were two isoforms of MAD1 detected by immunoblotting HeLa cell lysates, of which only the more slowly migrating form (MAD1α) co-immunoprecipitated with Cyclin B1 (Figure 1A). A previous study using hepatocellular carcinoma cells had identified an alternatively spliced form of MAD1 (MAD1β) (Sze et al., 2008) that lacks the 47amino acid-encoding exon 4 of MAD1α and migrates at the same molecular mass as our faster migrating form of MAD1. Therefore, we expressed MAD1α and MAD1β from cDNAs and found that these migrated at the same molecular masses as the two forms of MAD1 in cells, and that only MAD1α bound to Cyclin B1 (Figure S1C). In agreement with this, expressing a series of truncation mutants showed that residues 39 to 329 were able to bind to Cyclin B1 (data not shown).

**Figure 1.**
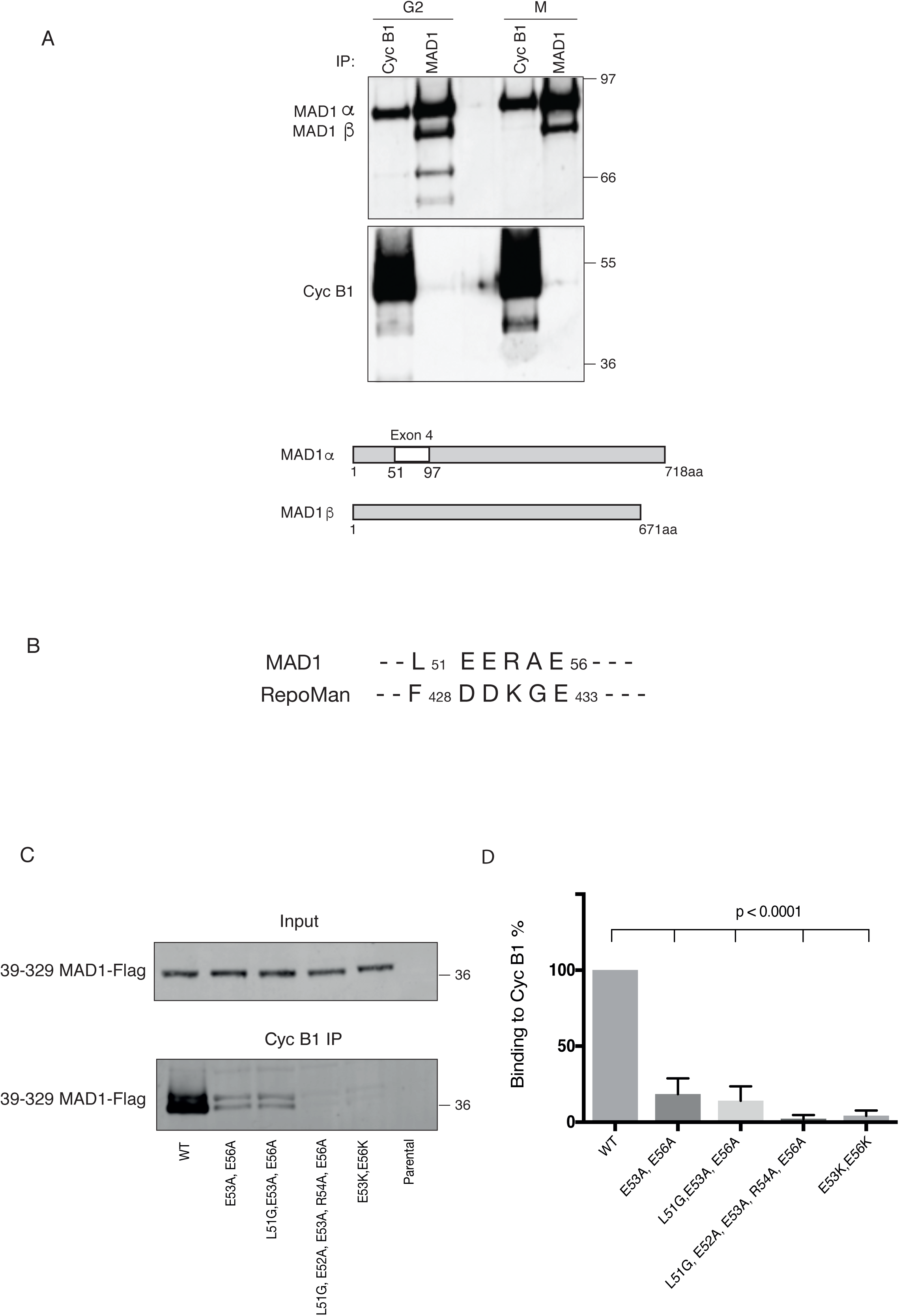
MAD1 binds to Cyclin B1 through the acidic face of a helix within exon 4. A) Hela cells were synchronised in either G2 phase or mitosis (M), Cyclin B1 was immunoprecipitated, subjected to SDS-PAGE and immunoblotted with anti-MAD1 (upper panel) and anti-Cyclin B1 antibodies (lower panel). Schematic shows the location of exon 4 in MAD1α that is absent from MAD1β. (B) Similarity between MAD1 and Repoman sequences within the regions found to interact with Cyclin B1. (C) Hela cells expressing wild-type or mutated MAD1-Flag (39-329aa) from a tetracyclin-inducible promoter were synchronised in either G2 phase or mitosis 12 hr after adding tetracyclin. Cyclin B1 was immunoprecipitated, subjected to SDS-PAGE, blotted with anti-FLAG antibody and assayed on a LiCOR Odyssey scanner. (D) The data from 3 experimental repeats were normalised to the amount of Cyclin B1 binding to wild type MAD1 and plotted using Prism software. Error bars represent SD, significance calculated using an unpaired t-test.

These analyses implicated the peptide sequence encoded by exon 4 (residues 51 to 97) as important for binding to Cyclin B1; however, exon 4 also contains the Nuclear Localisation Sequence (NLS) of MAD1, previously shown to be important for its function and its proper localisation to the nuclear pore complex in interphase (Sze et al., 2008). Therefore, we sought to narrow down the residues required to interact with Cyclin B1. A region of the RepoMan protein between amino acids 403-550 had been reported to bind Cyclin B1 (Qian 2015), and when we compared this region to exon 4 of MAD1 we found a small region of similarity (Figure 1B). This region of MAD1 is predicted to be part of helical region (using the JPred program, Cole et al., 2008), and likely to form a coiled-coil structure (McDonnell et al., 2006). An interaction with an acidic surface could conceivably be used to confer specificity for binding to B-type cyclins in animal cells because comparing the structures of B and A type cyclins shows that B-type cyclins are distinguished by their conserved basic patches at the interface with Cdk2 (Brown et al., 2007). Therefore, we mutated two acidic residues, E53 and E56, within the small region of MAD1 on the face of the helix not predicted to be involved in any coiled-coil interactions (Figure S1D). This double charge substitution of E53/56K severely perturbed binding to Cyclin B1 in vitro (Figure 1C) but not the localisation of full-length MAD1 to the nuclear envelope in interphase (Figure S1E).

### MAD1 recruits Cyclin B1 to kinetochores

We sought to identify the function of the binding between MAD1 and Cyclin B1. We used CRISPR/Cas9^D10A^ (Figure S2A) to introduce the E53/56K mutation into both alleles of MAD1 in RPE1 cells that had one allele of Cyclin B1 tagged with the Venus yellow fluorescent protein, and one allele of MAD2 tagged with the Ruby red fluorescent protein (Collin et al., 2013). We confirmed the MAD1 point mutations in two independent clones (7D2 and 8B12) by PCR analysis and genome sequencing (Figure S2B). In agreement with our in vitro pull downs, we found that the MAD1 E53/56K mutants were unable to bind Cyclin B1 (Figure 2A), but were still able to bind to MAD2 (Figure 2B) as expected since MAD2 binds to a region of MAD1 450 amino acids away. Mutating MAD1 in the RPE1 Cyclin B1-Venus:Ruby-MAD2 cells allowed us to assay Cyclin B1 and MAD2 recruitment to kinetochores in living cells. This showed that in both clones the E53/56K mutation prevented Cyclin B1 but not MAD2 from being recruited to unattached kinetochores (Figure 2C and movies S1A, S1B). We were thus in a position to determine the role of MAD1 binding to Cyclin B1 and its recruitment to kinetochores.

**Figure 2.**
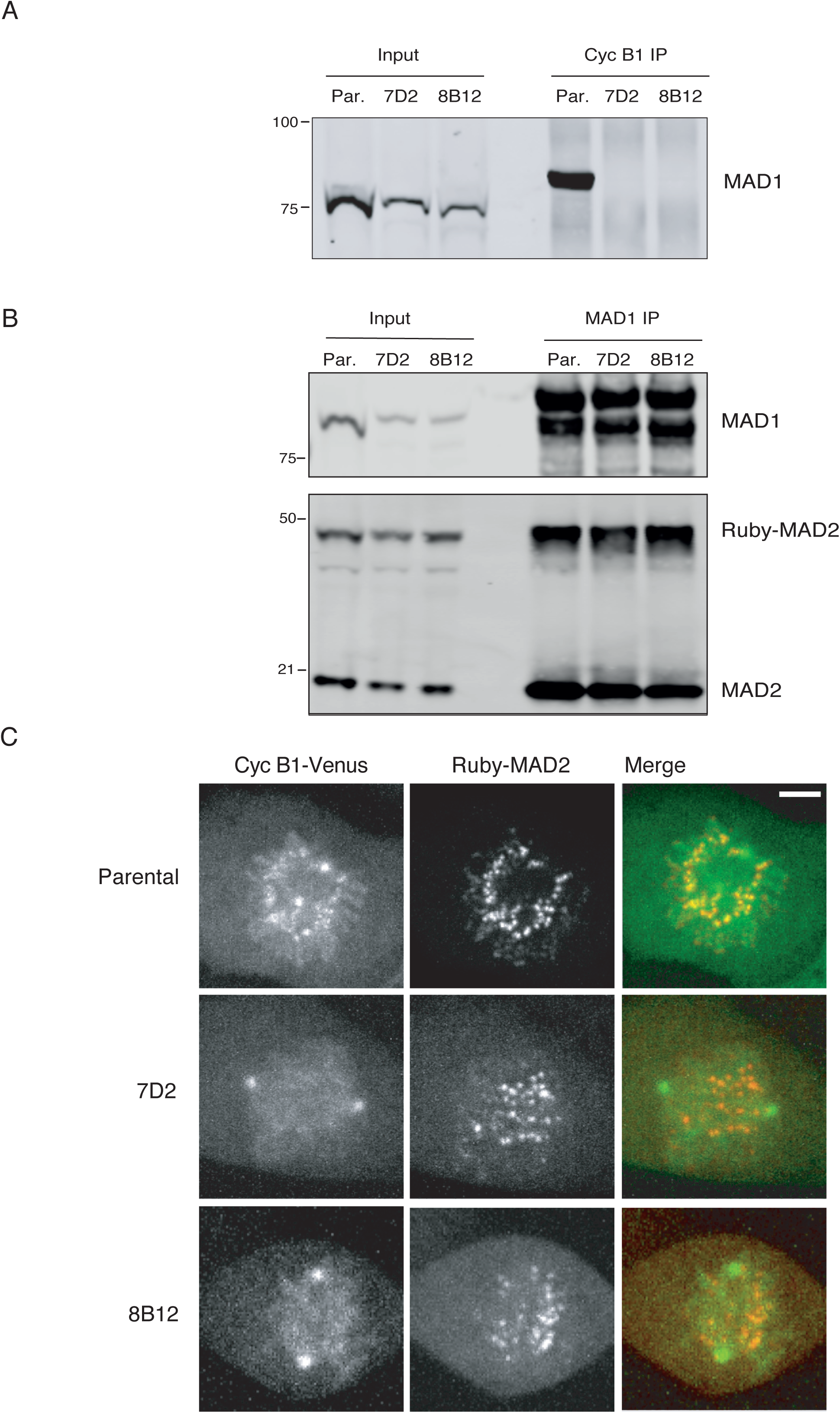
The E53K/E56K mutation prevents MAD1 binding to Cyclin B1 but not MAD2. (A) Parental RPE Cyclin B1-Venus^+/-^:Ruby-MAD2^+/-^ cells (Par.) or clones 7D2 and 8B12 carrying a homozygous mutation of E53/E56K in MAD1 were synchronised to enrich for G2 phase and mitosis, Cyclin B1 was immunoprecipitated, subjected to SDS-PAGE and immunoblotted with anti-MAD1 antibodies. (B) Parental RPE Cyclin B1-Venus^+/-^:Ruby-MAD2^+/-^ cells (Par.) or clones 7D2 and 8B12 were synchronised in G2 and M phase, MAD1 was immunoprecipitated, and immunoblotted with anti-MAD1 (upper panel) and anti-MAD2 (lower panel) antibodies. (C) Parental RPE Cyclin B1-Venus^+/-^:Ruby-MAD2^+/-^ cells or MAD1 E53/E56K clones 7D2 and 8B12 were assayed by spinning disk confocal time-lapse microscopy. Images of maximum intensity projections of Cyclin B1-Venus (left, green), Ruby-MAD2 (middle, red) and the merged image (right) for a representative prometaphase cell are shown, see movies S1A, S1B, S1C. Data shown for all panels are representative of 3 independent experiments. Scale bar = 3 *µ*M.

### MAD1 binding to Cyclin B1 is required for genomic stability

We first analysed the chromosomal stability of our MAD1 E53/56K clones compared to the parental RPE1 cells by counting the chromosome number in metaphase spreads. This showed that mutating MAD1 dramatically increased chromosomal instability: 77% (7D2) or 85% (8B12) of cells in the mutant clones had gained or (predominantly) lost chromosomes compared to less than 24% in the parental (Figure 3A). The increase in chromosomal instability in the MAD1 E53/56K clones might be explained by a weaker SAC. To test this, we assayed the SAC under three conditions: untreated cells, where the time from NEBD to anaphase is determined by the SAC (Figure 3B); mitotic delay in cells treated with low doses of nocodazole (Figure 3C); and mitotic delay in cells treated with paclitaxel (Figure 3D). There was no difference in timing between the wild type and mutant clones in low doses of nocodazole, but there was a significant difference between the parental and one mutant clone in unperturbed mitosis, and the other clone in response to paclitaxel. Paclitaxel treatment is a more sensitive assay for the SAC than nocodazole treatment since we have previously shown that fewer kinetochores recruit MAD2, and less MAD2 is recruited to these kinetochores, than in nocodazole (Collin et al., 2013). Thus, we conclude that MAD1 recruitment of Cyclin B1 to kinetochores strengthens the SAC and that this is important for long term genomic stability.

**Figure 3.**
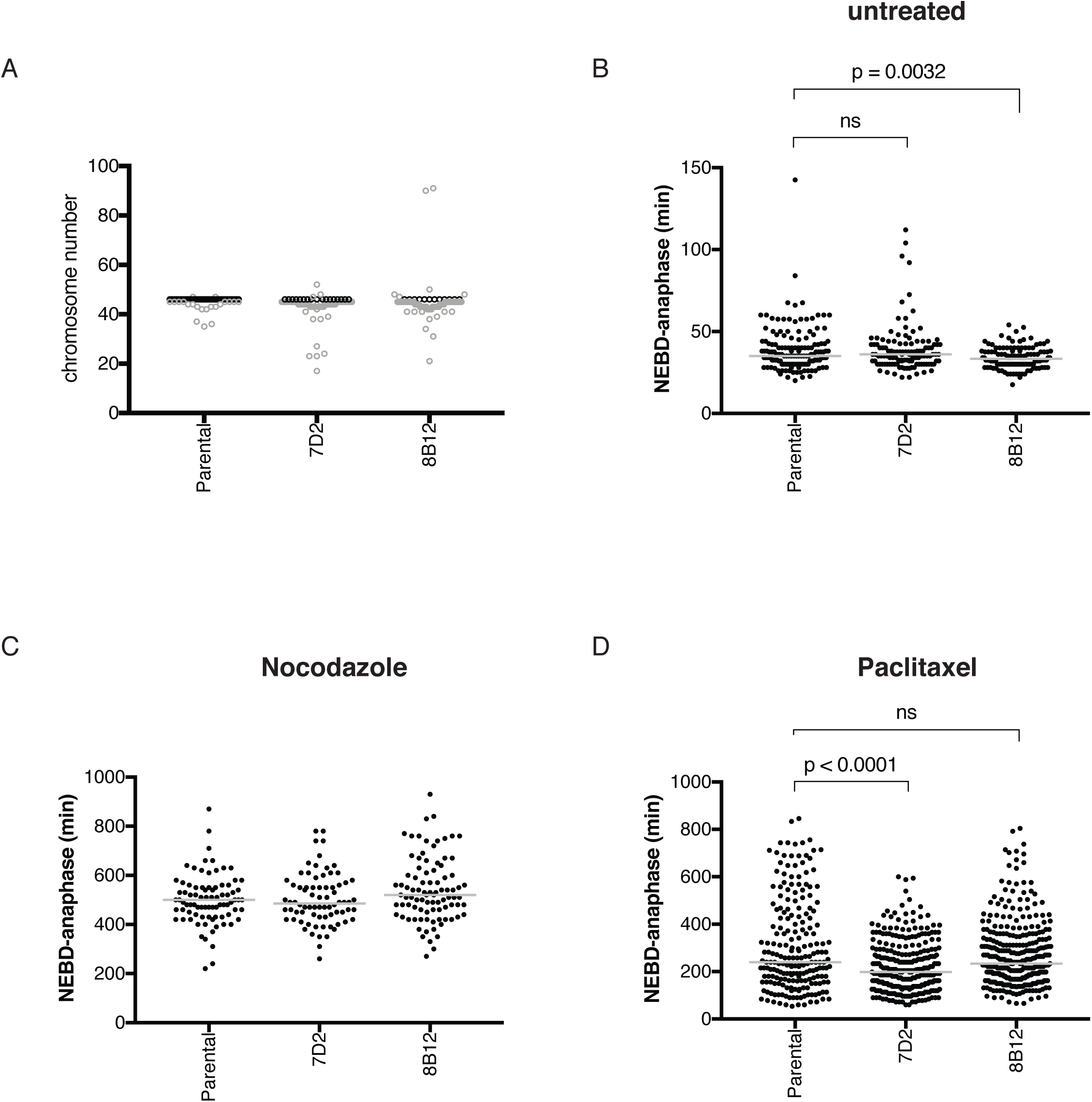
MAD1 binding to Cyclin B1 is required for genomic stability. (A) Chromosome number per cell was assayed for Parental RPE Cyclin B1-Venus^+/-^:Ruby-MAD2^+/-^ cells and the MAD1 E53/E56K clones 7D2 and 8B12 by metaphase spreads (n=89, n=74, n=80, respectively) in 3 independent experiments. Black dots indicate 46 chromosomes. (B) The time from NEBD to anaphase was measured for Parental RPE Cyclin B1-Venus^+/-^:Ruby-MAD2^+/-^ cells and the MAD1 E53/E56K clones 7D2 and 8B12 by time-lapse DIC microscopy. Scatter dot blots show the median time (grey bar) from three experiments (Parental= 147 cells, 7D2=119 cells, 8B12=126 cells). (C & D) The duration of the mitotic arrest for Parental RPE Cyclin B1-Venus^+/-^:Ruby-MAD2^+/-^ cells and the MAD1 E53/E56K clones 7D2 and 8B12, stained with sir-DNA, was assayed by time-lapse microscopy in 55nM nocodazole (Parental=83 cells, 7D2=76 cells, 8B12=90 cells in 3 experiments) (C) or 100 nM paclitaxel (Parental=210 cells, 7D2=296 cells, 8B12=325 cells in 3 experiments) (D). The median is shown as a grey line and p values in panels B and D were calculated using a Mann-Whitney unpaired t-test.

### Cells with MAD1 mutants that cannot bind Cyclin B1 are sensitive to partial inhibition of MPS1

In our previous studies on the strength of SAC signalling we showed that there was a strong inverse correlation between the strength of the SAC and the dose of an MPS1 inhibitor (Collin et al., 2013). We reasoned that partial inhibition of MPS1 might sensitise cells and uncover a more significant role for Cyclin B1-CDK1 binding to MAD1. In agreement with this, when we treated cells with low doses of an MPS1 inhibitor, either reversine (Fig 4) or AZ3146 (Figure S3), we found that the SAC was much more severely compromised in the MAD1 E53/56K mutant clones than in the parental cells, as assayed either by the timing from NEBD to anaphase (Figure 4A) or by the ability of cells to arrest in nocodazole (Figure 4B). Furthermore, live cell analyses of chromosome behaviour in these cells revealed that around 80% of the MAD1 mutant cells failed to form a proper metaphase plate and performed anaphase with a large number of lagging chromosomes, compared to less than 20% of the parental cells (Figure 4C and see movies S2A, S2B, S2C, S2D).

**Figure 4.**
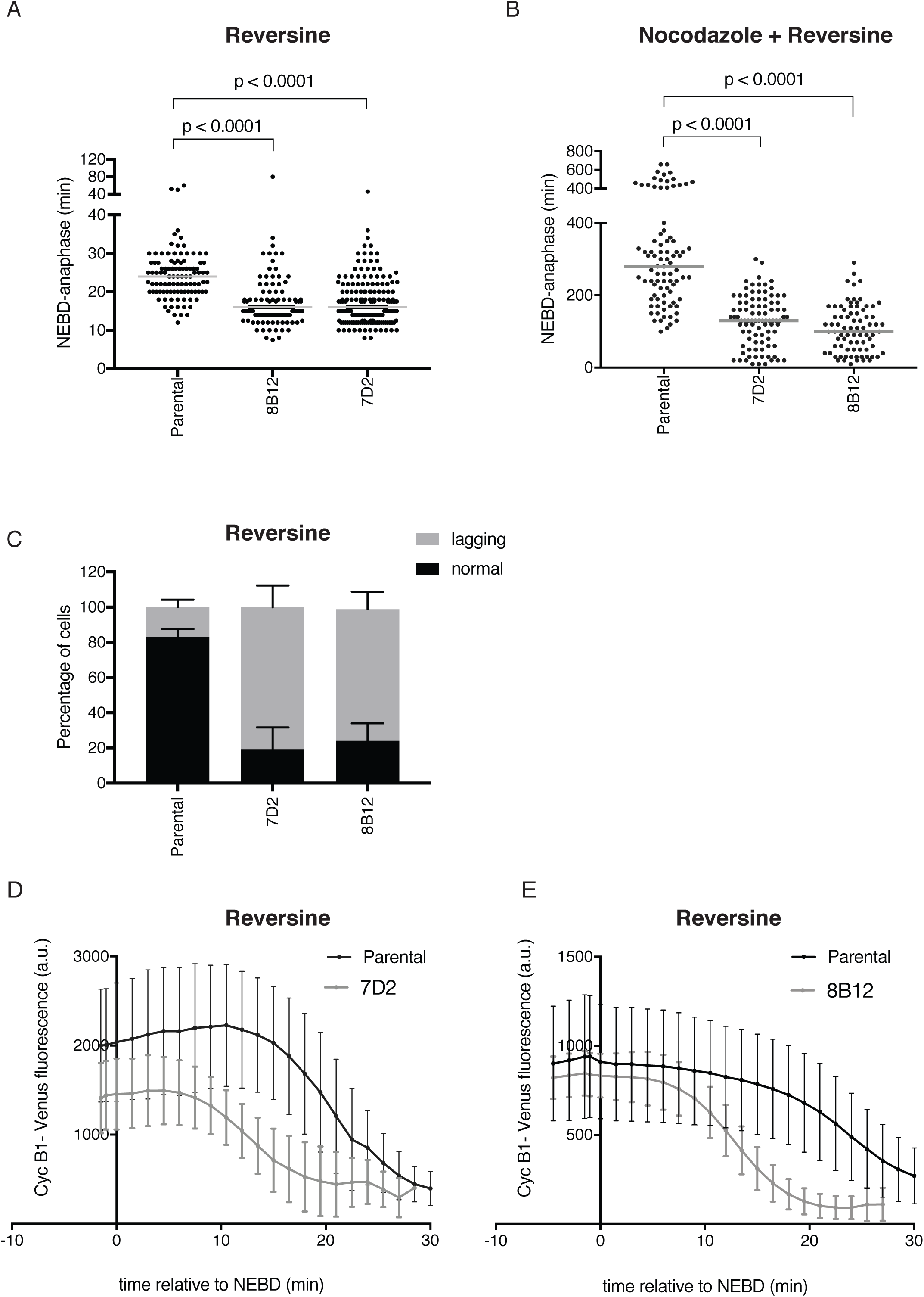
Cells with MAD1 mutants that cannot bind Cyclin B1 are sensitive to partial MPS1 inhibition. (A and B) The timing from nuclear envelope breakdown (NEBD) to anaphase was measured in Parental RPE Cyclin B1-Venus^+/-^:Ruby-MAD2^+/-^ cells and in the MAD1 E53/E56K clones 7D2 and 8B12. Cells were treated with 166nM reversine (Panel A, Parental= 104 cells, 7D2=189 cells, 8B12= 108 cells) or 55nM nocodazole plus 166nM reversine (Panel B, Parental= 86 cells, 7D2=93 cells, 8B12= 79 cells) and the data plotted as a scatter plot using Prism software. Median values shown as a grey line and the p values were calculated using an unpaired Mann Whitney t-test. Data from at least 3 independent experiments. (C) Quantification of lagging chromosomes in 166nM reversine-treated Parental RPE Cyclin B1-Venus+/-, Ruby-MAD2 +/-MAD1 wt and the MAD1 E53/E56K 7D2 and 8B12 clones stained with sir-DNA (see movies S2A and S2B). (D and E) Quantification of Cyclin B1-Venus degradation in Parental RPE Cyclin B1-Venus^+/-^:Ruby-MAD2^+/-^ cells and in the MAD1 E53/E56K clones 7D2 (D) and 8B12 (E) treated with 166 nM reversine. The total Cyclin B1-Venus fluorescence level in a cell was measured over time using time-lapse fluorescence microscopy. The data for 25 parental and 22 mutant cells (D) and for 23 parental and 35 mutant cells (E) are plotted as mean and SD. Data for 3 experimental repeats for panels D and E.

The premature sister chromatid separation exhibited by the MAD1 mutant cells could have one of two explanations: either they were unable to activate the SAC, or they were unable to maintain the SAC as the number of signalling kinetochores diminished following microtubule attachment. To distinguish between these two possibilities, we analysed the kinetics of Cyclin B1-Venus degradation (Figure 4D). Cyclin B1-Venus was stable for a few minutes after NEBD in the MAD1 mutant cells, but was then degraded with similar kinetics to wild type cells that have satisfied the SAC. This showed that the SAC was initially active in the mutant cells but it could not be maintained (Figure 4D and Figure 4E). Thus, we conclude that the SAC is much more dependent on MPS1 kinase activity when MAD1 is unable to recruit Cyclin B1.

### MAD2 recruitment to kinetochores is delayed when MAD1 cannot bind Cyclin B1

To gain insight into the mechanism underlying the weaker SAC signalling and the greater dependence on MPS1 in cells where MAD1 cannot bind Cyclin B1, we studied the recruitment of SAC proteins to kinetochores as cells began mitosis. To do this we used CRISPR/Cas9^D10A^ to introduce an RFP670 fluorescent tag into the MIS12 protein (Figures S4A, S4B and S4C, see also movies S2C and S2D) so that we could identify kinetochores in living cells. We then assayed the recruitment of MAD2 to kinetochores by quantifying the co-localisation between MAD2 and MIS12 in living cells (see Methods). This showed a striking difference between the parental and MAD1 E53/56K mutant clones (Figure 5A and 5B). In parental cells, MAD2 began to be recruited to the newly formed kinetochores 10 minutes or more before NEBD, whereas recruitment was markedly delayed in the MAD1 mutant cells; indeed, in the cells of one clone (7D2) MAD2 was not recruited until NEBD. In cells of the other clone (8B12), MAD2 recruitment was less delayed but much slower than normal and never reached the amounts seen in parental cells (Figure 5B). Thus, we conclude that kinetochores in MAD1 E53/56K mutant cells are compromised in their SAC signalling due to perturbed recruitment of MAD2.

**Figure 5.**
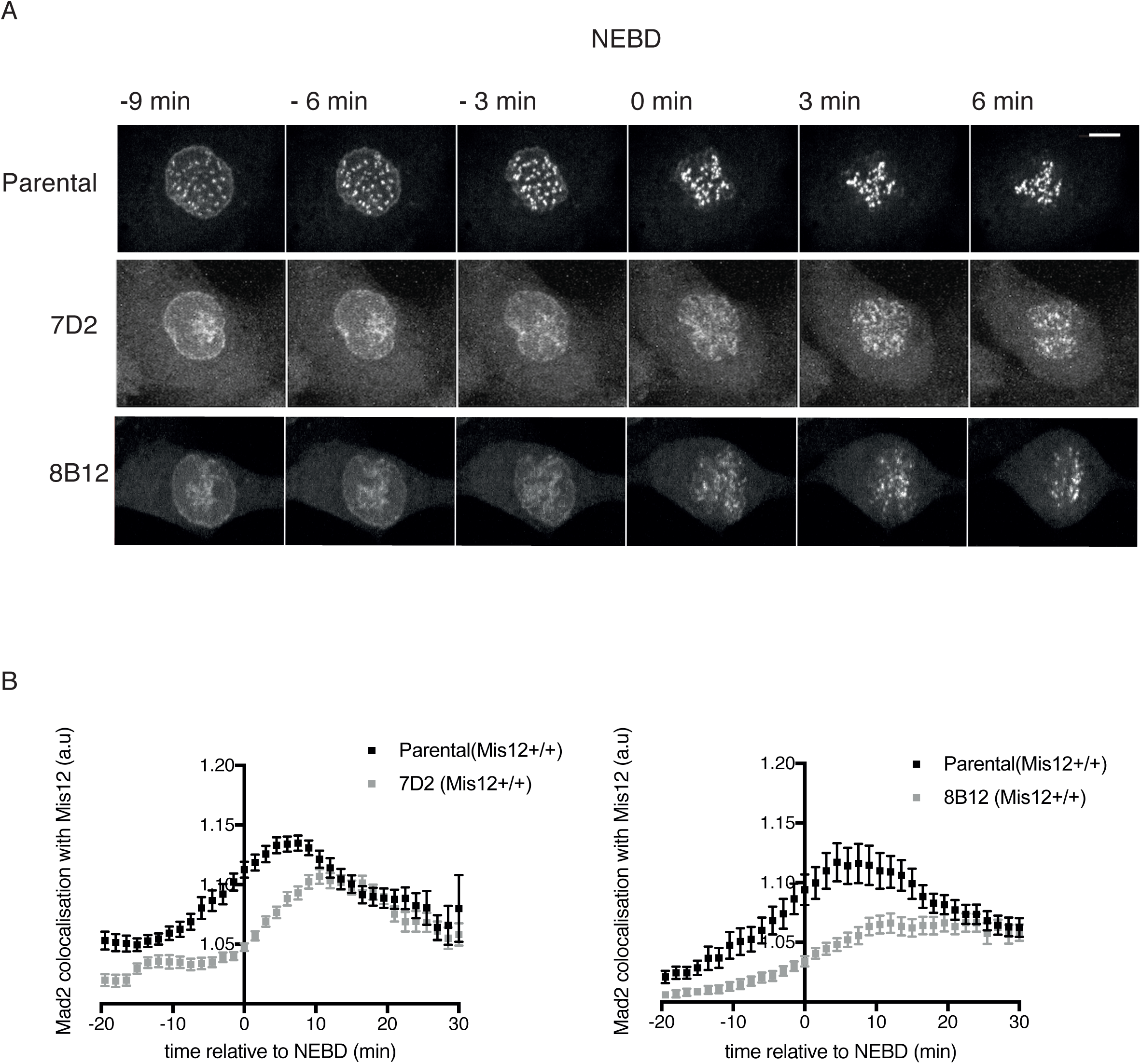
MAD2 recruitment to kinetochores is delayed when MAD1 cannot bind Cyclin B1. (A) Maximum intensity projections at the indicated times from time-lapse fluorescence of Parental RPE Cyclin B1-Venus^+/-^:Ruby-MAD2^+/-^ cells and the MAD1 E53/E56K clones 7D2 and 8B12 showing Ruby-MAD2 localisation relative to NEBD. Scale bar = 5µm. (B) Quantification of Ruby-MAD2 co-localisation with MIS12-RFP670 using the Coloc-3DT program (see Materials and Methods) relative to NEBD. Graphs show values obtained from at least 40 cells for each clone in 3 independent experiments, error bars indicate Standard Error of the Mean (SEM).

### MAD1 remains associated with TPR and condensing chromosomes when it cannot bind Cyclin B1

MAD2 has to bind MAD1 to be recruited to kinetochores (Chen et al., 1998); therefore, we analysed the behaviour of MAD1 in mutant and parental cells as they entered mitosis. We found that like MAD2, MAD1 recruitment to kinetochores was also perturbed when MAD1 was unable to bind Cyclin B1; this was because the MAD1 now appeared to associate with the condensing chromosomes (Figure 6A). We hypothesised that this might be caused by inefficient release from the nuclear basket; therefore, we analysed the behaviour of the TPR protein that binds MAD1 and is required for its localisation to the NPC (Lee et al., 2008). In agreement with our hypothesis, this revealed that mutant MAD1 that cannot bind Cyclin B1 remained associated with TPR on chromosomes in early mitosis (Figure 6A).

**Figure 6.**
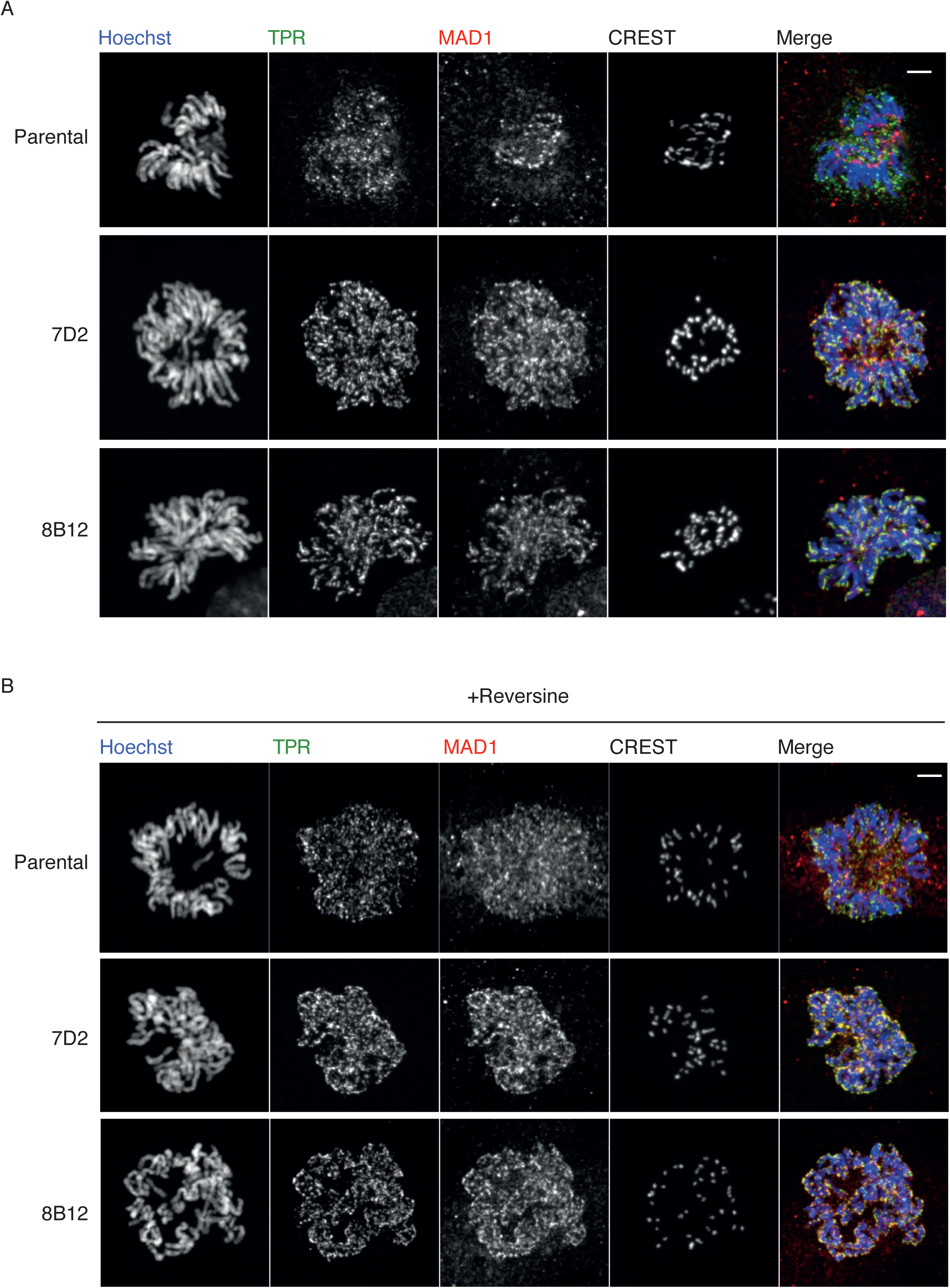
MAD1 remains associated with TPR and condensing chromosomes when it cannot bind Cyclin B1. (A) Prometaphase Parental RPE Cyclin B1-Venus^+/-^:Ruby-MAD2^+/-^ cells and the MAD1 E53/E56K clones 7D2 and 8B12 were fixed and stained with Hoechst 33342 (blue), anti-TPR (green), anti-MAD1 (Red) and CREST antibodies as indicated. (B) Cells were fixed and stained as in panel A except that they were pretreated with 166nM reversine. Scale bar = 2µm. Images are representative of 2 independent experiments.

We then asked whether the release of MAD1 from TPR might also be sensitive to MPS1 kinase activity and analysed the localisation of TPR and MAD1 in cells treated with a low dose of an MPS1 inhibitor. This had a striking effect in the MAD1 E53/56K cells where TPR and MAD1 almost completely co-localised around the chromosomes and very little MAD1 was able to bind to the newly formed kinetochores (Figure 6B). We observed a similar but much milder effect on MAD1 in Parental RPE cells (Figure 6B).

### The effect of the MAD1 mutant is partially rescued by directing Cyclin B1 to the NPC

Our results indicated that localised Cyclin B1-CDK1 kinase activity acts with MPS1 activity to release MAD1 from TPR and allow its recruitment to kinetochores; therefore, we postulated that targeting Cyclin B1 to the nuclear pore might rescue the effect of the MAD1 E53/56K mutant on the SAC. To test this, we tagged an mTurquoise2 (mTurq2)-labelled GFP-binding protein nanobody (GBP) to the C-terminus of the POM121 nuclear pore protein that binds the inner nuclear membrane (Hallberg et al., 1993), and randomly integrated the cDNA encoding this fusion protein into RPE1 parental Cyclin B1-Venus:Ruby-MAD2 cell line and into MAD1 E53/56K clone 7D2. Live cell imaging revealed that cells expressing the POM121-mTurq2-GBP fusion protein recruited Cyclin B1-Venus to the NPC (Figure 7A). We then treated these cells with 100nM paclitaxel plus 166nM reversine and assayed the ability of these cells to maintain a mitotic arrest. We compared the behaviour of these cells to cells expressing randomly integrated POM121 fused to mTurquoise2 alone, and the parental Cyclin B1-Venus:Ruby-MAD2 cells expressing either POM121 or POM121-GBP fusion proteins as controls (Figure 7B). In four separate experiments, we found that cells expressing the POM121-GBP fusion protein were able to maintain a mitotic arrest for significantly longer than cells expressing POM121. Thus, we conclude that local Cyclin B1-CDK1 activity at the NPC is responsible for the timely release of MAD1 from the NPC, which is required to generate a robust SAC and maintain genomic stability.

**Figure 7.**
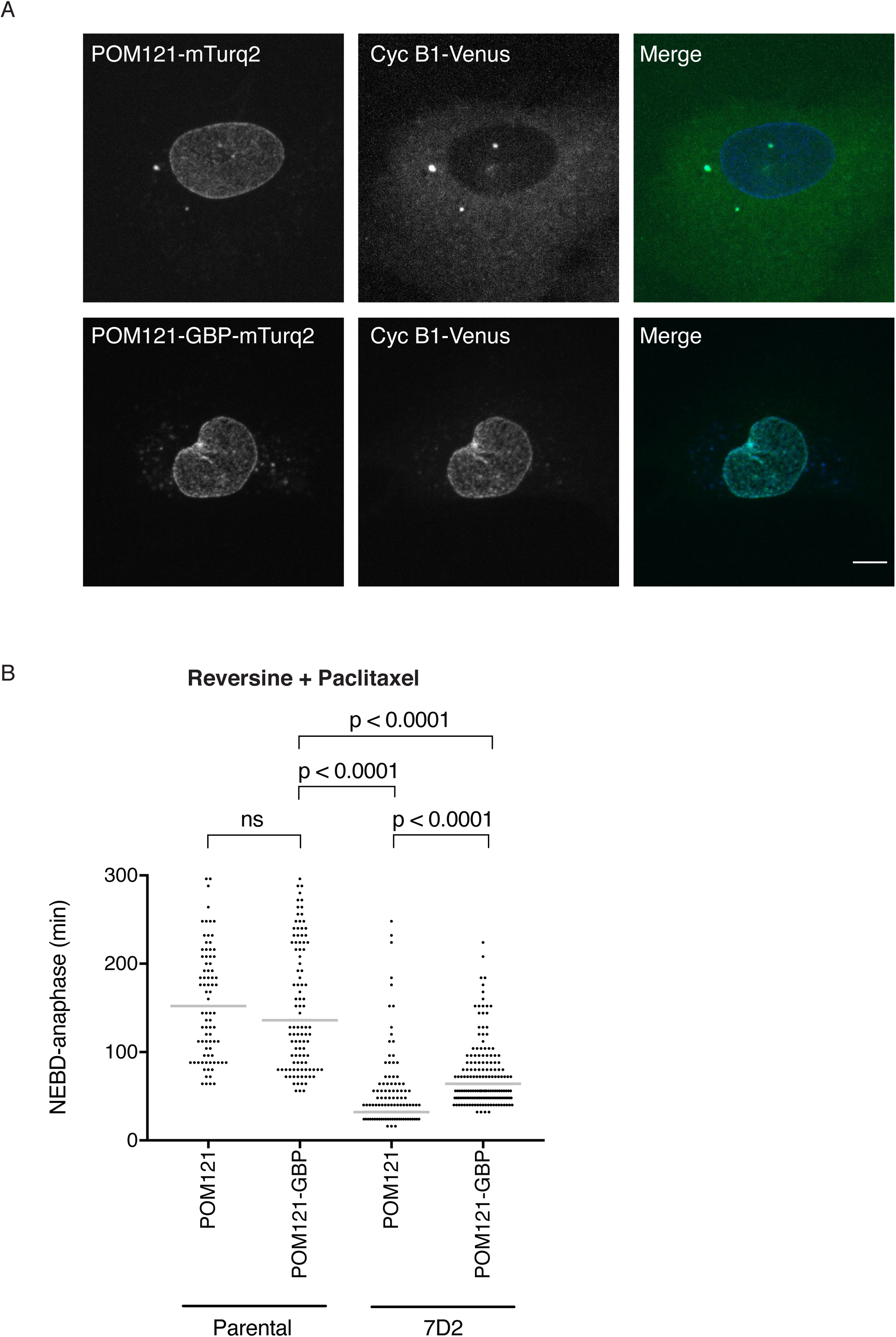
Targetting Cyclin B1 to the NPC partially restores the SAC. **A)** Maximum projection images of Parental RPE Cyclin B1-Venus^+/-^:Ruby-MAD2^+/-;^ RFP670-MIS12^+/+^ cells expressing POM121-mTurquoise2 (top panel) or POM121-GBP-mTurquoise2 (bottom panel). Left panels show the localisation of POM121-mTurq2 and POM121-GBP-mTurq2 at the NPC; middle panels show Cyclin B1-Venus; right panels are the merged images. Scale bar= 5µm (B) The duration of the mitotic arrest for Parental RPE Cyclin B1-Venus^+/-^:Ruby-MAD2^+/-^ cells and the MAD1 E53/E56K clone 7D2 expressing POM121-mTurquoise2 or POM121-GBP-mTurquoise2 were assayed by time-lapse microscopy in 166nM reversine and 100nM paclitaxel and the data plotted using Prism software. The p values are calculated using a Mann Whitney unpaired t-test. For the POM121-GBP-mTuquoise2-expressing cells, only cells where Cyclin B1-Venus was clearly recruited to the nuclear envelope were analysed.

## Discussion

In this study, we have shown that Cyclin B1 binds to the MAD1 protein through a predicted acidic patch on a helix of MAD1. Unexpectedly, we find that MAD1 recruits Cyclin B1-CDK1 to promote its own release from the NPC before NEBD, and that Cyclin B1-CDK1 coordinates with the MPS1 kinase to achieve this. The importance of releasing MAD1 before NEBD is that this allows MAD1 to bind to the newly formed kinetochores where it can begin to generate the MCC to inhibit the APC/C at NEBD. Thus, our findings identify a simple but elegant mechanism by which the rising level of Cyclin B1-CDK1 activity before NEBD both activates the APC/C and sets up the conditions to inhibit it until all the chromosomes attach to the mitotic spindle, thereby ensuring genomic stability.

We identified MAD1 as the most prominent interaction partner of Cyclin B1-CDK1. We, and others (Alfonso-Pérez et al., 2019) identified the N-terminus of MAD1 as the binding site for Cyclin B1. It is intriguing to note that this binding site is lost in the MAD1β spliced form that is prominent in hepatocellular carcinoma cell lines (Sze et al., 2008); it is conceivable that the inability to bind Cyclin B1, along with the loss of the nuclear localisation signal, might contribute to their genomic instability. We subsequently narrowed down the Cyclin B1 binding motif to a predicted acidic patch on a helical region of MAD1. Although beyond the scope of our present study, we are currently in the process of characterising this through structural studies, and determining whether this is a conserved interaction motif for other mitotic substrates of Cyclin B1. If so this will be, to our knowledge, the first interaction motif specific for the major mitotic kinase in animal cells. It is interesting to note that recognition of a helix may be a conserved feature of the cyclins since D-type cyclins recognise a predicted helix in the C-terminus of pRb (Topacio et al., 2019).

It is notable that preventing MAD1 from binding to Cyclin B1 perturbs its release from the NPC even though there is plenty of active Cyclin B1-CDK1 freely diffusing within the cell. Thus, our study identifies an important function for localised Cyclin B1-CDK1 activity and adds to our understanding of the increasing importance of local kinase-phosphatase gradients in controlling the cell (reviewed in Pines and Hagan, 2011). Spatial control of Cyclin B-CDK1 has been clearly demonstrated in triggering mitosis from the spindle pole body in fission yeast (Grallert et al., 2013; Hagan and Grallert, 2013) as has the spatial control of Plk1 through its recruitment to substrates previously phosphorylated by CDK1 (Elia et al., 2003a, 2003b), and in the control of error correction at kinetochores through the balance of Aurora B and PP1/PP2A phosphatases activities (Liu et al., 2010; Welburn et al., 2010; reviewed in Gelens et al., 2018; Liu et al., 2010).

In addition to emphasising the importance of spatial control of the mitotic kinases, our study also identifies how the reorganisation of the interphase cell is important for the subsequent function of mitosis-specific structures; in particular, how the disassembly of the NPC is coordinated with assembly of a functional kinetochore. The connection between NPC components and the kinetochore has been known for some time: in addition to MAD1/MAD2, the Nup107-160 complex, Nup358/RanBP2, and Crm1 proteins all associate with the kinetochores (Arnaoutov et al., 2005; Joseph et al., 2004; Zuccolo et al., 2007). Our findings now show how the timing of NPC disassembly is important for timely recruitment of MAD1 and MAD2 to kinetochores.

The role of CDK1 in NPC disassembly has been most clearly shown in studies using a powerful in vitro system (Linder et al., 2017; Marino et al., 2014). These studies have implicated the Plk1 and Nek kinases working in coordination with CDK1 to phosphorylate the core NPC components Nup98 and Nup53. Our study now reveals coordination between CDK1 and the MPS1 kinase in freeing MAD1 from TPR at the inner-NPC “basket”, and explains why only the initial localisation of MAD1 at kinetochores depends upon MPS1 activity. MAD1 localisation to unattached kinetochores is only abolished if MPS1 is inhibited before mitotic entry (Hewitt et al., 2010), i.e before release of MAD1 from the NPC.

It is intriguing to note that Plk1 and MPS1 recognise the same primary consensus motif (□-D/E-X-S) (Dou et al., 2011) and that they can cooperate by phosphorylating the same sites on proteins at the kinetochore (von Schubert et al., 2015). Therefore, it is tempting to speculate that some of the NPC components postulated to be phosphorylated by Plk1 might also be substrates of MPS1.

Once MAD1 has been released from TPR it binds to unattached kinetochores, where it can continue to recruit Cyclin B1. The binding of Cyclin B1 to unattached kinetochores has been observed by a number of groups (Alfonso-Pérez et al., 2019; Bentley et al., 2007; Chen et al., 2008), and this ‘guilt by association’ is one piece of evidence implicating Cyclin B1-CDK1 in the mechanics of the SAC and chromosome attachment. The problem in interpreting previous studies designed to elucidate the role of Cyclin B1-CDK1 in the SAC is the many SAC-independent roles that the kinase plays: it prevents cells from separating their sister chromatids and exiting mitosis; it maintains outer kinetochore structures; it prevents the activation of Cdh1; and it represses phosphatase activity (Holt et al., 2008; Qian et al., 2015; Visintin et al., 1998; Zachariae et al., 1998). Additional caveats are introduced in studies using small molecule inhibitors, which can affect other Cyclin-CDK1 family members and related kinase families. Thus, it has been difficult to ascribe a direct role for Cyclin B1-CDK1 in the SAC. We have overcome these problems by identifying and characterising a point mutant of MAD1 that prevents Cyclin B1 from being recruited to the kinetochore but leaves the rest of the Cyclin B1-CDK1 population active in the cell. A recent study used a large deletion mutant of MAD1 to address the same question but this mutant lacked 100 amino acids from the amino terminus of MAD1 (Alfonso-Pérez et al., 2019), thereby removing a number of other important functional domains, including the nuclear localisation signal that is required for it to bind to the NPC and the ability to form a stable putative coiled coil region that may contribute to kinetochore binding. This study concluded that by binding cyclin B1-CDK1, MAD1 increased MPS1 recruitment to kinetochores. We show here that MAD1 has first to bind Cyclin B1 to be efficiently released from the NPC and properly recruited to the kinetochore. MAD1 binding to Cyclin B1 could subsequently play a role at unattached kinetochores later in mitosis but this is beyond the scope of our present study. Moreover, our ability to strengthen the SAC in the MAD1 mutants by ectopically targeting Cyclin B1 to the NPC through POM121 shows that localised Cyclin B1-CDK1 activity is important for the proper control of mitosis.

Finally, our study reveals the mechanism by which the cell uses Cyclin B1-CDK1 to coordinate activation of the APC/C at NEBD with its immediate inhibition by the MCC to ensure genomic stability. We show here that Cyclin B1-CDK1 binding to MAD1 triggers MAD1/MAD2 release and recruitment to the newly formed kinetochore 10 minutes or more before NEBD and APC/C activation (den Elzen and Pines, 2001; Di Fiore and Pines, 2010; Geley et al., 2001), which should be sufficient time to generate a pool of MCC to inhibit the APC/C immediately upon NEBD. This model has the benefit that it simplifies the mechanisms required to inhibit the APC/C in early mitosis since the source of the MCC is the canonical unattached kinetochore. Its importance is underlined by the genomic instability manifested when MAD1 can no longer bind to Cyclin B1.

## Materials and Methods

### Plasmids and cell lines

MAD1 was tagged at the C-terminus with a 3xHA-Flag epitope by PCR and cloned into a modified version of pcDNA5 FRT/TO (ThermoFisher Scientific). Full length MAD1 carrying either L51G/E52A/E53A/R54A/E56 or E53Q/E56Q mutations were tagged at the C-terminus with mRuby by sub-cloning into the pMCSV vector. The POM121 coding region was amplified by PCR from POM121-EGFP_3_ plasmid (kind gift from Martin Hetzer, Salk Institute) and tagged at the C-terminus by sub-cloning into a modified version of pMCSV containing wither mTurquoise2 or GBP (GFP-binding protein)-mTurquoise2. To generate stable cell lines, parental RPE1 and clones 7D2 and 8B12 all expressing Cyclin B1-Venus:Ruby-MAD2:RFP670-MIS12 were transfected with POM121-mTurquoise2 and POM121-GBP-mTurquoise2 and cells were selected with 0.4 μg/mL neomycin (GIBCO). All constructs were verified by sequencing and sequences are available on request.

### Cell culture and Synchronisation

HeLa FRT/TO and RPE1 cells were cultured as described (Mansfeld et al., 2011). HeLa FRT/TO cells were transfected using the Flp-in-System (ThermoFisher Scientific). Cells were induced with tetracycline (1ug/ml; Calbiochem) 12 hours before harvesting. HeLa FRT/TO cells were synchronised in S phase by a double thymidine (2.5mM) block, then either released for 10 hours for G2 phase arrested extracts, or for mitotic cells, released into nocodazole (0.33mM) for 14 hours before mitotic cells were collected by shake off. RPE1 cells were synchronised in G2 phase through a 24-hour treatment with 100 nM palbociclib (Selleckchem) followed by 14 hours release into fresh medium.

### Drug treatments

For live-cell experiments, cells were treated with 50 nM sir-DNA (Spirochrome) for 3 hours before filming. AZ3146 (0.62 μM, Tocris), Paclitaxel (100 nM, Sigma-Aldrich), Reversine (166 nM, Cambridge BioScience), Nocodazole (55 nM, Sigma-Aldrich) were added just prior to filming.

### Genome editing

Genome editing was performed using CRISPR/Cas9^D10A^ technology. For the MAD1 E53/56K mutation, a donor plasmid (pJ241-305516 MAD1 E53K/E56K, synthesised by ATUM, California) comprising 12 silent point mutations in addition to the E53/56K substitutions and flanked by 400bp (5’) and 800bp (3’) sequences, was linearised with NotI digestion. The linearised plasmid was purified (GeneJET Gel Extraction Kit, ThermoFisher Scientific), and co-transfected into RPE1 Cyclin B1-Venus:Ruby-MAD2 (Collin et al., 2013)together with a modified version of the PX466 “All-in-One” plasmid (Chiang et al., 2016)containing Cas9D10A-T2A-RFP670 and gRNAs targeting MAD1 exon4 (5’-TCACTGAGGATTCTGTTTTT-3’ and 5’-GGTGCGACCTGCTCAGCTGG-3’). RFP670-expressing cells were selected using a FACSAria™ III (BD Biosciences) and sorted individually into a 96-well plate. For genotyping, genomic DNA was prepared using DirectCell-PCR Lysis-Reagent Cell (VWR) according to the manufacturer’s protocol and screened by PCR using a FailSafe™ PCR kit (Buffer E, Epicentre). The presence of MAD1 E53/56K substitutions was identified through PCR using forward primers annealing to the mutated or the wild type sequences (AGCTGGAAAAGAGGGCGAAAC and TAAGTGCCGGGAGATGCTG, respectively) and the same reverse primer (AGCCCACACAACGCACACCGA). Positive clones for the E53/56K mutations were screened using primers annealing ∼200bp upstream and downstream of the point mutations. PCR products were separated on agarose gels, cloned into the pDrive vector (Qiagen) and sequenced as shown in Fig. S2. The MIS12 locus was targeted with RFP670 as shown in Supplementary Fig.4. A donor plasmid containing RFP670 sequence in frame with MIS12 exon1 flanked by homology regions was co-transfected with the “All-in-One” plasmid comprising MIS12 specific gRNAs (5’-ATGACCTACGAGGCCCAGTT-3’ and 5’-CGCCACAAACGTGCATGCTT-3’) and Cas9D10A-T2A-EGFP. EGFP-positive cells were selected via FACS then sorted individually by FACS 10 days later. RFP670 positive clones were identified by PCR (as shown in Fig. S4B) and subsequently analysed by live-cell microscopy to confirm MIS12 expression and localisation (Fig. S4C).

### Metaphase spreads

Cells were treated with 0.1 μg/ml colcemide (GIBCO) for 3 hours, trypsinized, washed twice with 1X PBS and resuspended in hypotonic buffer (0.075 M potassium chloride). After a 20 minute incubation at 37°C, cells were centrifuged and the pellet was gently resuspended in 3:1 methanol/glacial acetic acid fixative (vortex dropwise). Cells were washed with fixative 3 times and a few drops were released onto an alcohol cleaned slide and allowed to air dry. Slides were counterstained with KaryoMAX™ Giemsa Stain Solution (GIBCO). Transmitted light images of metaphase spreads were captured using a 63x 1.4NA lens and the number of chromosomes per cell was counted using ImageJ software.

### Immunoprecipitation

Cells were lysed in lysis buffer (0.5% NP40 w/v 140 mM NaCl, 10mM KCl, 50 mM Hepes pH 7.2, 10% w/v glycerol, 1mM EDTA, HALT protease inhibitor cocktail (ThermoFisher Scientific). Supernatants from 11.000 x g centrifugation of cell lysates were incubated with anti-Cyclin B1 (GNS1, Pharminogen) or anti-MAD1 (9B10, Sigma-Aldrich) antibodies coupled to Protein G-Dynabeads (ThermoFisher Scientific) for 1 hour at 4°C, and washed four times in lysis buffer, and eluted at 65°C for 5 min before analysis by SDS-PAGE and silver or Colloidal Blue staining, or immunoblotting. Silver staining was performed according to manufacturers’ instructions (SilverQuest, Sigma-Aldrich) and Colloidal Blue staining as previously described (Rowley et al., 2000).

### Mass Spectrometry (MS)

For MS analyses immunoprecipitates on Protein-G Dybabeads were washed 2x with TEAB Buffer (100mM) and incubated with Trypsin (Roche) at 37°C for 18 hours. The tryptic peptides were collected and TMT-labelled according to manufacturers’ instructions (ThermoFisher Scientific). The TMT peptides were fractionated on a U3000 HPLC system (ThermoFisher Scientific) using an XBridge BEH C18 column (2.1 mm id × 15 cm, 130 Å, 3.5 *µ*m, Waters) at pH 10, with a 30min linear gradient from 5 - 35% acetonitrile (ACN)/NH_4_OH at a flow rate at 200 *µ*l/min. The fractions were collected every 30sec into a 96-wellplate by rows, then concatenated by columns to 12 pooled fractions and dried in a SpeedVac. The peptides were re-dissolved in 0.5% formic acid (FA) before LC-MS/MS analysis. The LC-MS/MS analysis were performed on the Orbitrap Fusion Lumos mass spectrometer coupled with U3000 RSLCnano UHPLC system (ThermoFisher Scientific). The peptides were first loaded to a PepMap C18 trap (100 *µ*m i.d. x 20 mm, 100 Å, 5 *µ*m) for 8 min at 10 *µ*l/min with 0.1% FA/H_2_O, then separated on a PepMap C18 column (75 *µ*m i.d. x 500 mm, 100 Å, 2 *µ*m) at 300 nl/min and a linear gradient of 8-30.4% ACN/0.1% FA in 120 min /cycle at 150 min for each fraction. The data acquisition used the SPS5-MS3 method with Top Speed at 3s per cycle time. The full MS scans (m/z 375-1500) were acquired at 120,000 resolution at m/z 200, and the AGC was set at 4e5 with 50ms maximum injection time. The most abundant multiply-charge ions (z = 2-5, above 10,000 counts) were subjected to MS/MS fragmentation by CID (35% CE) and detected in an ion trap for peptide identification. The isolation window by quadrupole was set m/z 0.7, and AGC at 10,000 with 50ms maximum injection time. The dynamic exclusion window was set ±7 ppm with a duration at 40sec, and only single charge status per precursor was fragmented. Following each MS2, the 5-notch MS3 was performed on the top 5 most abundant fragments isolated by Synchronous Precursor Selection (SPS). The precursors were fragmented by HCD at 65% CE then detected in Orbitrap at m/z 100-500 with 50,000 resolution for peptide quantification data. The AGC was set 100,000 with maximum injection time at 105ms.

### Data Analysis

The LC-MS/MS data were processed in Proteome Discoverer 2.2 (ThermoFisher Scientific) using SequestHT and Mascot search engines against the SwissProt protein database (v. August 2018) plus cRAP contaminant database (ftp://ftp.thegpm.org/fasta/cRAP). The precursor mass tolerance was set at 15 ppm and the fragment ion mass tolerance was set at 0.5 Da. Spectra were searched for fully tryptic peptides with maximum of two miscleavages. TMT6plex (Peptide N-terminus, K) was set as static modification, and dynamic modifications included Deamidation (N, Q), Oxidation (M) and Phosphorylation (S,T,Y). Peptides were validated by Percolator with q value threshold set at 0.05 for the decoy database search. Phosphorylation site locations were verified by the *ptmRS* module. The search result was filtered to achieve a protein FDR of 0.05. The TMT10plex reporter ion quantifier used 20 ppm integration tolerance on the most confident centroid peak at the MS3 level. Only unique peptides were used for quantification. Co-isolation threshold was set to 100%. Peptides with average reported S/N >3 were used for protein quantification. Only master proteins were reported. Only proteins with quantification values in all samples were used for further analyses. Protein abundances were normalised to the bait protein in each immunoprecipitation subset. To filter-out non-specific proteins, a LIMMA-based differential analysis was performed comparing MAD1 immunoprecipitations among themselves or versus IgG control samples. Proteins were deemed significantly different if adjusted p < 0.05 and two-fold difference in abundance.

### Immunoblotting

Immunoblotting was performed as previously described (Di Fiore and Pines, 2010). Primary antibodies were used at the indicated concentrations: MAD1 (9B10, Sigma 2mg/ml) 1/400; FLAG (M2, Sigma; 3mg/ml) 1/1000; MAD2 (Bethyl Laboratories cat A300-301A; 1mg/ml) 1/1000. IRDye680 and IRDye800CW (LI-COR)-conjugated secondary antibodies were used at 1:10,000. The signal was detected using an Odyssey scanner (LI-COR) as previously described (Nilsson et al., 2008).

### Immunofluorescence

Cells were fixed for 15 min at room temperature in PHEM (60mM Pipes, 25mM Hepes, 10mM EGTA, 2mM MgCl_2_ pH6.9 buffered with KOH) buffer with 4% w/v paraformaldehyde and 0.5% v/v TX100. Primary antibodies were diluted as follows: TPR (Atlas antibodies) 1/50; MAD1 (9B10, Sigma 2mg/ml) 1/200 and (Bethyl Laboratories cat no. A300-339) 1/500; Human Nuclear ANA-centromere autoantibody-Crest (Cortex Biochem) 1/200. Secondary antibodies were: anti-mouse-594nm, anti-rabbit-488nm, anti-human-647nm (Alexa Fluor, ThermoFisher Scientific), all at 1/400. Confocal imaging of antibody stained samples was performed on a Mariannis microscope (Intelligent Imaging Innovations, USA).

### Time-Lapse Imaging and Analysis

For time-lapse microscopy, cells were seeded and transfected on an 8-well chamber slide (μslide, Ibidi). Cells were pre-treated with 50 nM Sir-DNA (Spirochrome) 3hr before filming to visualise chromosomes. Cells were imaged in Leibovitz’s L-15 medium (GIBCO) supplemented with 10% FBS. Time-lapse confocal imaging was performed on a Marianas confocal spinning-disk microscope system (Intelligent Imaging Innovations, Inc., USA) comprising: a laser stack for 445 nm/488 nm/514 nm/561 nm lasers; an Observer Z1 inverted microscope (Carl Zeiss, Germany) equipped with Plan-Apochromat 40x 1.3NA and 63x 1.4 NA lenses; an OKO stage top incubator (OKO, Italy); a CSU X1 spinning disk head (Yokogawa, Japan); a Gemini W view optical splitter attached to a Flash4 CMOS camera (Hamamatsu, Japan) and a QuantEM 512SC camera (Photometrics, USA). The microscope was equipped with Brightline filters (Semrock, USA) for GFP/RFP, for CFP/YFP/RFP, and for RFP670. Immunofluorescene images were captured on a similar Marianas confocal microscope but equipped with a CSU W1 head. Colocalisation of Ruby-MAD2 with RFP670-MIS12 and immunofluorescence images were collected using a 63× 1.2 NA water corrected objective (Carl Zeiss, Germany). Time-lapse widefield fluorescence and DIC imaging was performed on a Nikon Eclipse microscope (Nikon, Japan) equipped with 20× 0.75 NA, 40× 1.3 NA and 63× 1.4 NA lenses, a Flash 4.0 CMOS camera (Hamamatsu, Japan), an excitation and an emission filter wheel equipped with Brightline (Semrock, USA) filters for CFP, GFP, YFP, RFP and RFP670 and an analyser in the emission wheel for DIC imaging. Image acquisition and processing for the confocal microscopes was performed using Slidebook 6 (Intelligent Imaging Innovation, Inc.) software; Micromanager software and ImageJ open source software were used for widefield imaging.

3D movies of Ruby-MAD2 localisation with RFP670-MIS12 of single cells were quantified using an open source program (https://github.com/adamltyson/coloc-3DT). Images were converted to OME-TIFF(Linkert et al., 2010), loaded into a custom python program, resliced in Z to isotropic sampling and smoothed with a Gaussian filter (sigma = 1 voxel). To segment the kinetochores, the MIS12-RF670 signal was thresholded using an adaptation of Otsu’s method (Otsu, 1979)in which the threshold was scaled by a fixed value (1.08) for all experiments. Noise was removed by morphological opening (kernel = 1 voxel cube), and then the mean value (colocalisation) of MAD2-Ruby was calculated within the thresholded kinetochores. This colocalisation was scaled to the level of Ruby-MAD2 within the rest of the nucleus (estimated as between 1-15 voxels from the segmented kinetochore). All image processing was performed with Scikit-image (van der Walt et al., 2014).

### Statistics

Statistical analyses were performed using GraphPad Prism. Significance of data derived from mitotic timings was determined using unpaired two-tailed Mann-Whitney tests. Statistical analyses of Mass Spectrometry were performed using the R package LIMMA and protein filtered for a logFC >2-fold and adjusted pValue <0.05. Binding to Cyclin B1 was analysed by unpaired two-tailed Student’s *t* tests.

## Author Contributions

MJ identified MAD1 binding to cyclin B1 the E53/56 MAD1 residues necessary for binding to Cyclin B1. MJ and CM generated and characterised the MAD1 E53/56K mutants. MJ, CM, MP, LY and JC performed the mass spectrometry, CM analysed the data. AT and MJ designed and wrote the coloc-3DT program. JP and CM designed and executed the POM121 rescue experiment. MJ, CM and JP analysed the data and wrote the paper.

## Acknowledgements

We thank; Stephen Taylor for the HeLa FRT cell line; Will Chiang and Steve Jackson for the CRISPR-Cas9^D10A^ construct; Martin Hetzer for POM121 cDNA; Marco Chiapello for LIMMA analysis; Andy Riddell, Fredrik Wahlberg, and Rhadika Patel for help with fluorescence activated cell sorting; Oxana Nashchekina for help with designing the strategy to mutate MAD1 in RPE1 cells; Federica Schiavoni for advice on metaphase spreads; Iain Hagan and Eleanor Trotter for the original idea and advice on synchronising cells with Palbociclib, and all the Cell Division Team for helpful discussions.

## Supplementary Figures and Movies Legends

### Supplementary Figures

**Figure S1. Related to Figure 1.**
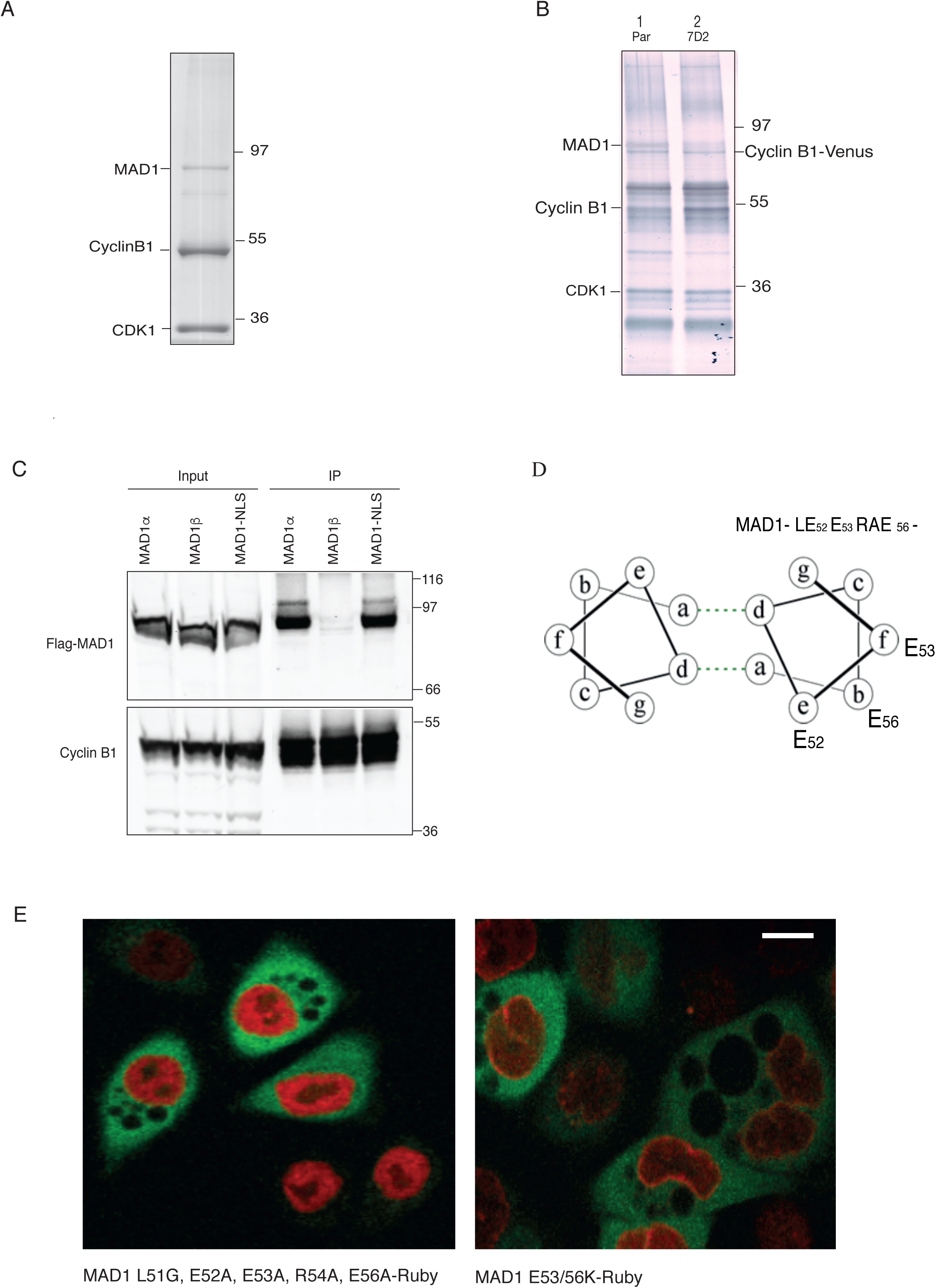
Cyclin B1 binds to MAD1 through the acidic face of a helix encoded by exon 4. (A). Colloidal blue stained SDS-PAGE gel of Cyclin B1 immunoprecipitated from HeLa cells. Marked bands were excised and identified by mass spectrometry. (B) Silver stained SDS-PAGE gel of Cyclin B1 immunoprecipitates from RPE Cyclin B1-Venus^+/-^: Ruby-MAD2^+/-^ cells (Lane 1) and MAD1 E53K/E56K clone 7D2 (Lane 2). (C) Cyclin B1 immunoprecipitates from Hela cells expressing Flag-epitope tagged MAD1α or MAD1β or MAD1-NLS KKR79-82AAA, probed with anti-FLAG (upper panel) or anti-Cyclin B1 (lower panel) antibodies. (D) Heptad registration of acidic residues of MAD1 within coiled-coil configuration, predicted using http://cb.csail.mit.edu/cb/paircoil2/ (McDonnell et al. 2006). (E) Confocal image of Hela Cyclin B1-Venus^+/-^ (green) cells transfected with either MAD1 L51G/E52A/E53A/R54A/E56A-Ruby (left panel, red) or MAD1 E53/56K-Ruby (right panel, red). Scale bar = 10*µ*M.

**Figure S2. Related to Figure 2.**
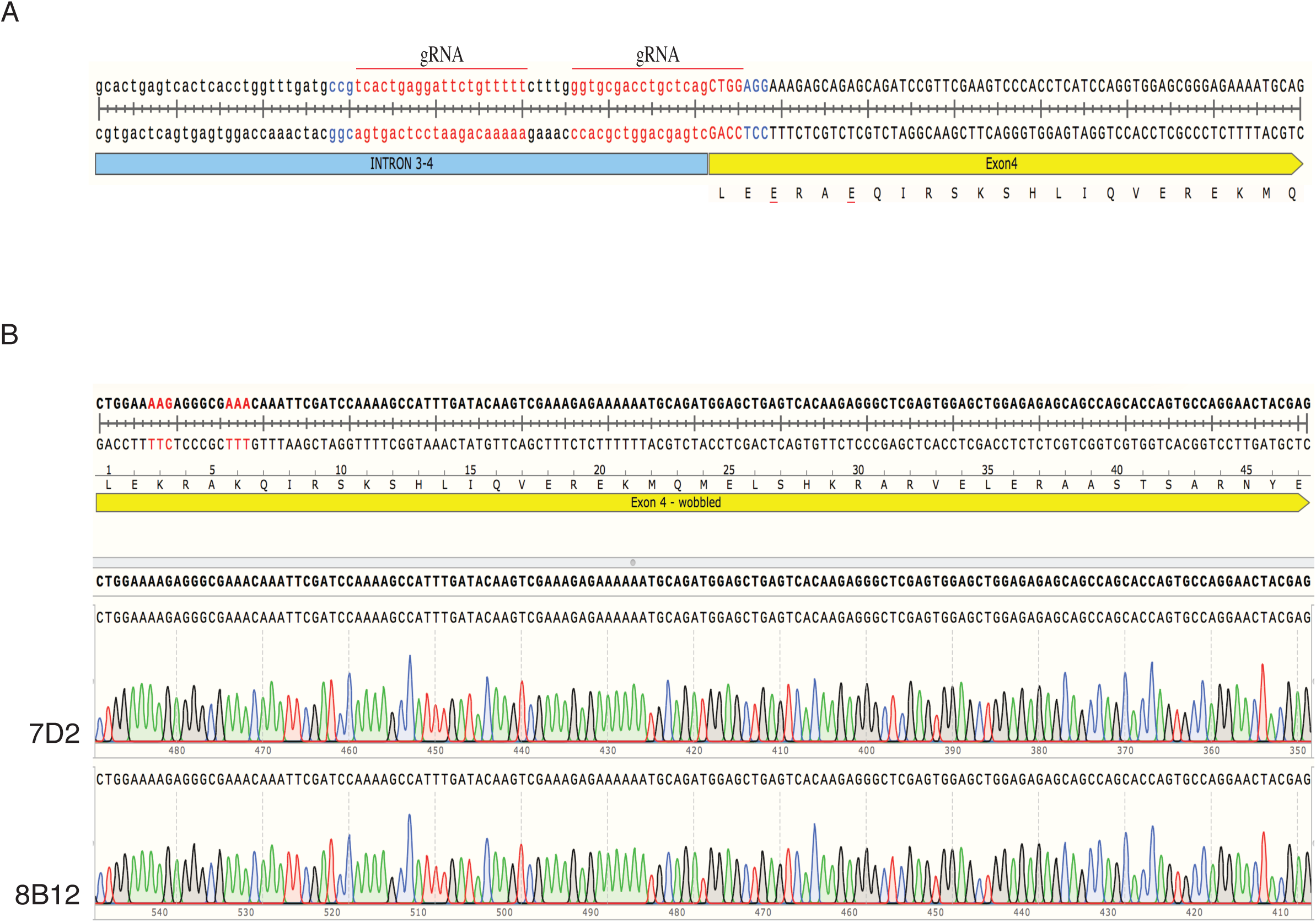
MAD1 recruits Cyclin B1 to kinetochores. (A) Schematic showing guide RNA (gRNA, red) selection and PAM (blue) for CRISPR-Cas9^D10A^ targeting of MAD1 exon4. (B) Genomic DNA sequencing of RPE Cyclin B1-YFP^+/-^ :MAD2-Ruby^+/-^ MAD1 E53/56K clones 7D2 and 8B12.

**Figure S3. Related to Figure 4.**
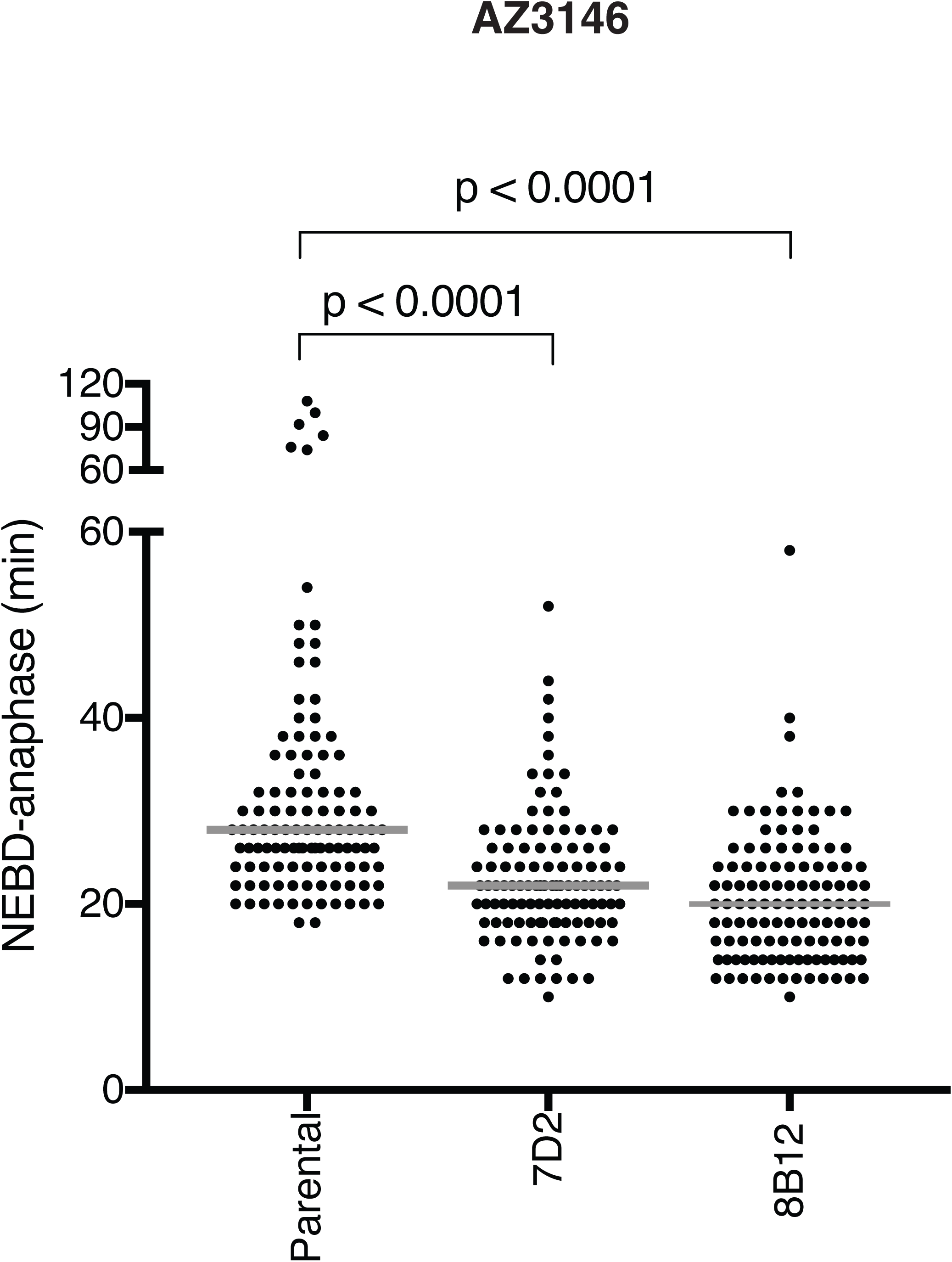
Cells with MAD1 mutants that cannot bind Cyclin B1 are sensitive to partial inhibition of MPS1. **(A)** Time from NEBD-anaphase for parental RPE Cyclin B1-YFP^+/-^:MAD2-Ruby^+/-^ cells and MAD1 E53/56K clones 7D2 and 8B12 treated with 0.62 μM AZ3146 MPS1 kinase inhibitor. Scatter dot blots show the median (grey line) from 2 independent experiments (Parental=112 cells, 7D2=111 cells, 8B12=117 cells).

**Figure S4. Related to Figure 5.**
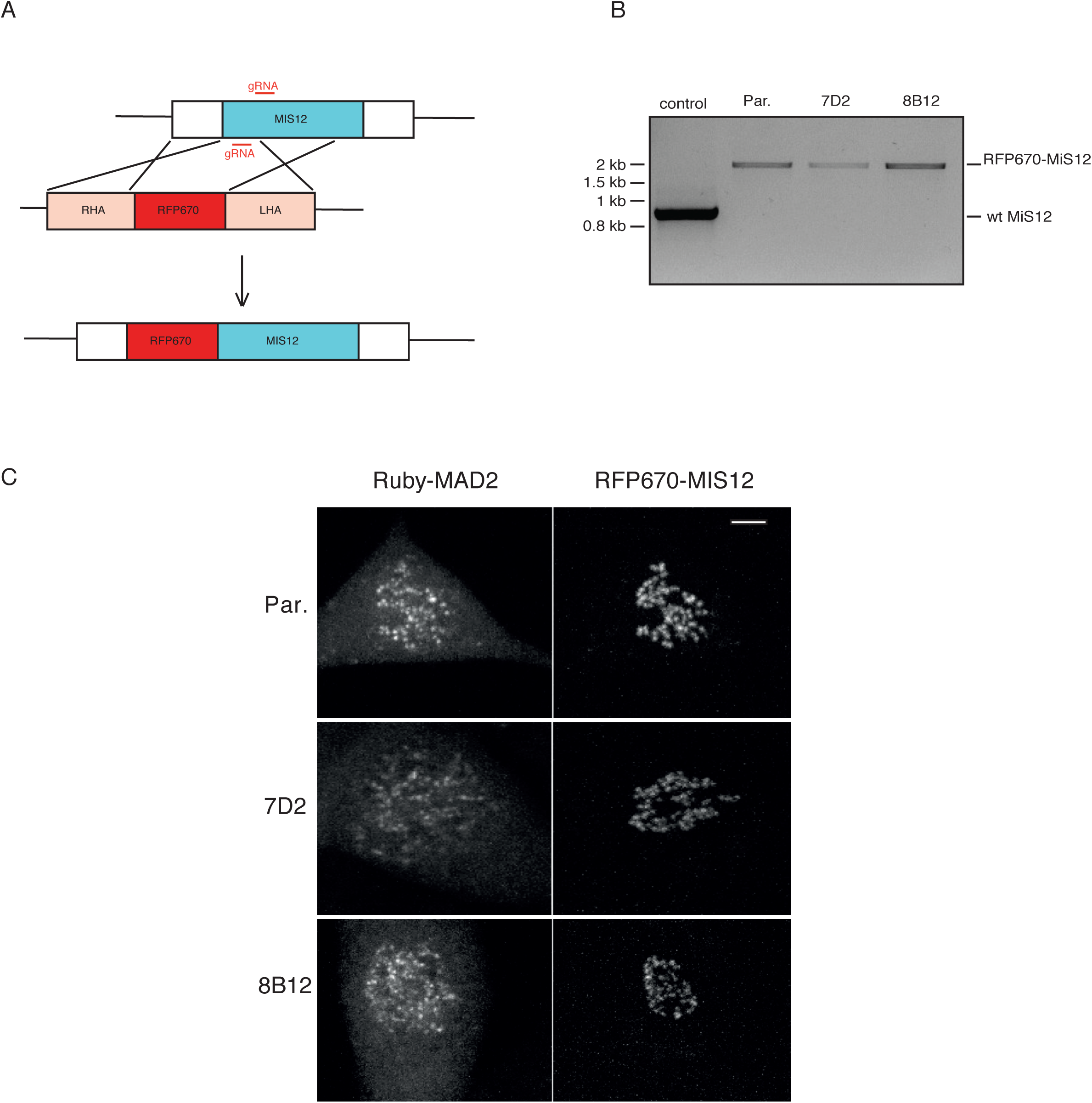
MAD2 recruitment to kinetochores is delayed when MAD1 cannot bind Cyclin B1. (A) Schematic showing how RFP670 was tagged at the amino-terminus of MIS12 (RHA and LHA refers to right and left homology arms, respectively). (B) PCR of genomic DNA from wild type RPE1 cells (control), Parental RPE Cyclin B1-Venus^+/-^: Ruby-MAD2^+/-;^ RFP670-MIS12+/+ cells (Par.) or MAD1 E53/E56K: RFP670-MIS12+/+ clones 7D2 and 8B12, showing integration of RFP670 into both alleles of MIS12. (C) Maximum projection images of Parental RPE Cyclin B1-Venus^+/-^:Ruby-MAD2^+/-;^ RFP670-MIS12^+/+^ cells (Par.) and MAD1 E53/E56K: RFP670-MIS12+/+ clones 7D2 and 8B12. Left panels show Ruby-MAD2; right panels show RFP670-MIS12. Scale bar, top right panel = 5*µ*m

### Supplementary Movies

Movie S1. Related to Figure 2 and S2. **MAD1 recruits Cyclin B1 to kinetochores**

Movies show mitotic entry of Parental RPE Cyclin B1-YFP^+/-^:MAD2-Ruby^+/-^ cell (movie S1A) or MAD1 E53/E56K clones 7D2 (movie S1B) and 8B12 cells (movie S1C); Cyclin B1-Venus (left panel), Ruby-MAD2 (middle panel) and merged channels Cyclin B1-Venus (green), Ruby-MAD2 (red) (right panel). Cells imaged by spinning disk confocal microscopy. Frame rate 1 image every 2 min.

Movies S2. Related to Figure. 4 **Cells with MAD1 mutants that cannot bind Cyclin B1 are sensitive to partial inhibition of MPS1**

Widefield epifluorescence movies show mitotic entry of Parental RPE Cyclin B1-Venus^+/-^ :Ruby-MAD2^+/-;^cells (S2A) or MAD1E53/E56K clones 8B12 (S2B) treated with sir-DNA 3 hours before filming and reversine (166nM) just before imaging. Frame rate 1 image every 2 min. (S2C&D) Spinning disk confocal movies show mitotic entry of (S2C) Parental RPE Cyclin B1-Venus^+/-^:Ruby-MAD2^+/-;^ RFP670-MIS12^+/+^ cells or (S2D) MAD1 E53/E56K clones 7D2. Cyclin B1-Venus (far left panel), Ruby-MAD2 (left panel, green), RFP-670-MIS12 (right panel, red) and merged channels for MAD2 and MIS12 (far right panel). Cells treated with reversine (166nM) just before imaging. Note that reversine reduces the loading of Cyclin B1 onto kinetochores in the parental cells. Frame rate 1 image every 2 min.

## Bibliography

Alexander, J., Lim, D., Joughin, B.A., Hegemann, B., Hutchins, J.R., Ehrenberger, T., Ivins, F., Sessa, F., Hudecz, O., Nigg, E.A., et al. (2011). Spatial Exclusivity Combined with Positive and Negative Selection of Phosphorylation Motifs Is the Basis for Context-Dependent Mitotic Signaling. Sci Signal 4, ra42, 1–15.

Alfonso-Pérez, T., Hayward, D., Holder, J., Gruneberg, U., and Barr, F.A. (2019). MAD1-dependent recruitment of CDK1-CCNB1 to kinetochores promotes spindle checkpoint signaling. J Cell Biol 218(4), 1108–1117.

Arion, D., Meijer, L., Brizuela, L., and Beach, D. (1988). cdc2 is a component of the M phase-specific histone H1 kinase: Evidence for identity with MPF. Cell 55, 371–378.

Arnaoutov, A., Azuma, Y., Ribbeck, K., Joseph, J., Boyarchuk, Y., Karpova, T., McNally, J., and Dasso, M. (2005). Crm1 is a mitotic effector of Ran-GTP in somatic cells. Nat Cell Biol 7, 626–632.

Bentley, A.M., Normand, G., Hoyt, J., and King, R.W. (2007). Distinct Sequence Elements of Cyclin B1 Promote Localization to Chromatin, Centrosomes, and Kinetochores during Mitosis. Mol Biol Cell 18, 4847–4858.

Bondt, H.L., Rosenblatt, J., Jancarik, J., Jones, H.D., Morgant, D.O., and Kim, S.-H. (1993). Crystal structure of cyclin-dependent kinase 2. Nature 363, 595–602.

Brandeis, M., Rosewell, I., Carrington, M., Crompton, T., Jacobs, M., Kirk, J., Gannon, J., and Hunt, T. (1998). Cyclin B2-null mice develop normally and are fertile whereas cyclin B1-null mice die in utero. Proc National Acad Sci 95, 4344–4349.

Brown, N.R., Noble, M.E., Endicott, J.A., and Johnson, L.N. (1999). The structural basis for specificity of substrate and recruitment peptides for cyclin-dependent kinases. Nat Cell Biol 1, 438–443.

Brown, N.R., Lowe, E.D., Petri, E., Skamnaki, V., Antrobus, R., and Johnson, L. (2007). Cyclin B and Cyclin A Confer Different Substrate Recognition Properties on CDK2. Cell Cycle 6, 1350–1359.

Cairo, L.V., Ptak, C., and Wozniak, R.W. (2013). Mitosis-Specific Regulation of Nuclear Transport by the Spindle Assembly Checkpoint Protein Mad1p. Mol Cell 49, 109–120.

Castilho, P.V., Williams, B.C., Mochida, S., Zhao, Y., and Goldberg, M.L. (2009). The M Phase Kinase Greatwall (Gwl) Promotes Inactivation of PP2A/B55δ, a Phosphatase Directed Against CDK Phosphosites. Mol Biol Cell 20, 4777–4789.

Champion, L., Linder, M.I., and Kutay, U. (2017). Cellular Reorganization during Mitotic Entry. Trends Cell Biol 27, 26–41.

Chen, Q., Zhang, X., Jiang, Q., Clarke, P.R., and Zhang, C. (2008). Cyclin B1 is localized to unattached kinetochores and contributes to efficient microtubule attachment and proper chromosome alignment during mitosis. Cell Res 18, 268–280.

Chen, R. H., Shevchenko, A., Mann, M., and Murray, A.W. (1998). Spindle Checkpoint Protein Xmad1 Recruits Xmad2 to Unattached Kinetochores. J Cell Biology 143, 283–295.

Cole, C., Barber, J.D., and Barton, G.J. (2008). The Jpred 3 secondary structure prediction server. Nucleic Acids Res 36, 197–201.

Collin, P., Nashchekina, O., Walker, R., and Pines, J. (2013). The spindle assembly checkpoint works like a rheostat rather than a toggle switch. Nat Cell Biol 15, 1378–1385.

D’Angiolella, V., Mari, C., Nocera, D., Rametti, L., and Grieco, D. (2003). The spindle checkpoint requires cyclin-dependent kinase activity. Gene Dev 17, 2520–2525.

Dasso, M. (2006). Ran at kinetochores. Biochem Soc T 34, 711–715.

den Elzen, N., and Pines, J. (2001). Cyclin a Is Destroyed in Prometaphase and Can Delay Chromosome Alignment and Anaphase. J Cell Biology 153, 121–136.

Di Fiore, B., and Pines, J. (2007). Emi1 is needed to couple DNA replication with mitosis but does not regulate activation of the mitotic APC/C. J Cell Biology 177, 425–437.

Di Fiore, B., and Pines, J. (2010). How cyclin A destruction escapes the spindle assembly checkpoint. J Cell Biology 190, 501–509.

Dorée, M., and Hunt, T. (2002). From Cdc2 to Cdk1: when did the cell cycle kinase join its cyclin partner? J Cell Sci 115, 2461–2464.

Dou, Z., von Schubert, C., Körner, R., Santamaria, A., Elowe, S., and Nigg, E.A. (2011). Quantitative Mass Spectrometry Analysis Reveals Similar Substrate Consensus Motif for Human Mps1 Kinase and Plk1. Plos One 6(4), e18793.

Dowdy, S.F., Hinds, P.W., Louie, K., Reed, S.I., Arnold, A., and Weinberg, R.A. (1993). Physical interaction of the retinoblastoma protein with human D cyclins. Cell 73, 499–511.

Dunphy, W.G., and Newport, J.W. (1989). Fission yeast p13 blocks mitotic activation and tyrosine dephosphorylation of the Xenopus cdc2 protein kinase. Cell 58, 181–191.

Elia, A., Rellos, P., Haire, L.F., Chao, J.W., Ivins, F.J., Hoepker, K., Mohammad, D., Cantley, L.C., Smerdon, S.J., and Yaffe, M.B. (2003a). The Molecular Basis for Phosphodependent Substrate Targeting and Regulation of Plks by the Polo-Box Domain. Cell 115, 83–95.

Elia, A.E., Cantley, L.C., and Yaffe, M.B. (2003b). Proteomic Screen Finds pSer/pThr-Binding Domain Localizing Plk1 to Mitotic Substrates. Science 299, 1228–1231.

Forbes, D.J., Travesa, A., Nord, M.S., and Bernis, C. (2015). Nuclear transport factors: global regulation of mitosis. Curr Opin Cell Biol 35, 78–90.

Frye, J.J., Brown, N.G., Petzold, G., Watson, E.R., Grace, C.R., Nourse, A., Jarvis, M.A., Kriwacki, R.W., Peters, J.-M., Stark, H., et al. (2013). Electron microscopy structure of human APC/CCDH1–EMI1 reveals multimodal mechanism of E3 ligase shutdown. Nat Struct Mol Biology 20, 827–835.

Fujimitsu, K., Grimaldi, M., and Yamano, H. (2016). Cyclin-dependent kinase 1–dependent activation of APC/C ubiquitin ligase. Science 352, 1121–1124.

Gascoigne, K.E., and Cheeseman, I.M. (2013). CDK-dependent phosphorylation and nuclear exclusion coordinately control kinetochore assembly state. J Cell Biology 201, 23–32.

Gavet, O., and Pines, J. (2010). Progressive Activation of CyclinB1-Cdk1 Coordinates Entry to Mitosis. Dev Cell 18, 533–543.

Gelens, L., Qian, J., Bollen, M., and Saurin, A.T. (2018). The Importance of Kinase–Phosphatase Integration: Lessons from Mitosis. Trends Cell Biol 28, 6–21.

Geley, S., Kramer, E., Gieffers, C., Gannon, J., Peters, J.-M., and Hunt, T. (2001). Anaphase-Promoting Complex/Cyclosome–Dependent Proteolysis of Human Cyclin a Starts at the Beginning of Mitosis and Is Not Subject to the Spindle Assembly Checkpoint. J Cell Biology 153, 137–148.

Gharbi-Ayachi, A., Labbé, J.-C., Burgess, A., Vigneron, S., Strub, J.-M., Brioudes, E., Van-Dorsselaer, A., Castro, A., and Lorca, T. (2010). The Substrate of Greatwall Kinase, Arpp19, Controls Mitosis by Inhibiting Protein Phosphatase 2A. Science 330, 1673–1677.

Golan, A., Yudkovsky, Y., and Hershko, A. (2002). The Cyclin-Ubiquitin Ligase Activity of Cyclosome/APC Is Jointly Activated by Protein Kinases Cdk1-Cyclin B and Plk. J Biol Chem 277, 15552–15557.

Grallert, A., Patel, A., Tallada, V.A., Chan, K., Bagley, S., Krapp, A., Simanis, V., and Hagan, I.M. (2013). Centrosomal MPF triggers the mitotic and morphogenetic switches of fission yeast. Nat Cell Biol 15, 88–95.

Hagan, I.M., and Grallert, A. (2013). Spatial control of mitotic commitment in fission yeast. Biochem Soc T 41, 1766–1771.

Hagting, A., Jackman, M., Simpson, K., and Pines, J. (1999). Translocation of cyclin B1 to the nucleus at prophase requires a phosphorylation-dependent nuclear import signal. Curr Biol 9, 680–689.

Hallberg, E., Wozniak, R.W., and Blobel, G. (1993). An integral membrane protein of the pore membrane domain of the nuclear envelope contains a nucleoporin-like region. J. Cell Biol. 122, 513–521. doi:10.1083/jcb.122.3.513

Hayward, D., Alfonso-Pérez, T., Cundell, M.J., Hopkins, M., Holder, J., Bancroft, J., Hutter, L.H., Novak, B., Barr, F.A., and Gruneberg, U. (2019). CDK1-CCNB1 creates a spindle checkpoint–permissive state by enabling MPS1 kinetochore localization. J. Cell Biol. 218(4), 1182–1199.

Heald, R., and McKeon, F. (1990). Mutations of phosphorylation sites in lamin A that prevent nuclear lamina disassembly in mitosis. Cell 61, 579–589.

Hein, J.B., and Nilsson, J. (2016). Interphase APC/C–Cdc20 inhibition by cyclin A2–Cdk2 ensures efficient mitotic entry. Nat Commun 7, 10975.

Hewitt, L., Tighe, A., Santaguida, S., White, A.M., Jones, C.D., Musacchio, A., Green, S., and Taylor, S.S. (2010). Sustained Mps1 activity is required in mitosis to recruit O-Mad2 to the Mad1–C-Mad2 core complex. J Cell Biology 190, 25–34.

Hirano, T. (2012). Condensins: universal organizers of chromosomes with diverse functions. Gene Dev 26, 1659–1678.

Holt, L.J., Krutchinsky, A.N., and Morgan, D.O. (2008). Positive feedback sharpens the anaphase switch. Nature 454, 353.

Jackman, M., Lindon, C., Nigg, E.A., and Pines, J. (2003). Active cyclin B1–Cdk1 first appears on centrosomes in prophase. Nat Cell Biol 5, 143–148.

Jeffrey, P.D., Russo, A.A., Polyak, K., Gibbs, E., Hurwitz, J., Massagué, J., and Pavletich, N.P. (1995). Mechanism of CDK activation revealed by the structure of a cyclinA-CDK2 complex. Nature 376, 313–320.

Joseph, J., Liu, S.-T., Jablonski, S.A., Yen, T.J., and Dasso, M. (2004). The RanGAP1-RanBP2 Complex Is Essential for Microtubule-Kinetochore Interactions In Vivo. Curr Biol 14, 611–617.

Labbe, J., Lee, M., Nurse, P., Picard, A., and Doree, M. (1988). Activation at M-phase of a protein kinase encoded by a starfish homologue of the cell cycle control gene cdc2+. Nature 335, 251–254.

Labit, H., Fujimitsu, K., Bayin, S.N., Takaki, T., Gannon, J., and Yamano, H. (2012). Dephosphorylation of Cdc20 is required for its C-box-dependent activation of the APC/C. Embo J 31, 3351–3362.

Lara-Gonzalez, P., Westhorpe, F.G., and Taylor, S.S. (2012). The Spindle Assembly Checkpoint. Curr Biol 22, 966–980.

Lee, S., Sterling, H., Burlingame, A., and McCormick, F. (2008). Tpr directly binds to Mad1 and Mad2 and is important for the Mad1–Mad2-mediated mitotic spindle checkpoint. Gene Dev 22, 2926–2931.

Linder, M.I., Köhler, M., Boersema, P., Weberruss, M., Wandke, C., Marino, J., Ashiono, C., Picotti, P., Antonin, W., and Kutay, U. (2017). Mitotic Disassembly of Nuclear Pore Complexes Involves CDK1- and PLK1-Mediated Phosphorylation of Key Interconnecting Nucleoporins. Dev Cell 43, 141–156.

Liu, D., Vleugel, M., Backer, C.B., Hori, T., Fukagawa, T., Cheeseman, I.M., and Lampson, M.A. (2010). Regulated targeting of protein phosphatase 1 to the outer kinetochore by KNL1 opposes Aurora B kinase. J Cell Biology 188, 809–820.

Loïodice, I., Alves, A., Rabut, G., van Overbeek, M., Ellenberg, J., Sibarita, J.-B., and Doye, V. (2004). The Entire Nup107-160 Complex, Including Three New Members, Is Targeted as One Entity to Kinetochores in Mitosis. Mol Biol Cell 15, 3333–3344.

London, N., and Biggins, S. (2014). Mad1 kinetochore recruitment by Mps1-mediated phosphorylation of Bub1 signals the spindle checkpoint. Gene Dev 28, 140–152.

Lu, D., Hsiao, J.Y., Davey, N.E., Voorhis, V.A., Foster, S.A., Tang, C., and Morgan, D.O. (2014). Multiple mechanisms determine the order of APC/C substrate degradation in mitosis. J Cell Biol 207, 23–39.

Mansfeld, J., Collin, P., Collins, M.O., Choudhary, J.S., and Pines, J. (2011). APC15 drives the turnover of MCC-CDC20 to make the spindle assembly checkpoint responsive to kinetochore attachment. Nat Cell Biol 13, 1234–1243.

Marino, J., Champion, L., Wandke, C., Horvath, P., Mayr, M.I., and Kutay, U. (2014). Chapter 12 An In Vitro System to Study Nuclear Envelope Breakdown. Methods Cell Biol 122, 255–276.

McDonnell, A., Jiang, T., Keating, A., and Berger, B. (2006). Paircoil2: improved prediction of coiled coils from sequence. Bioinformatics 22, 356–358.

McGrath, D.A., Balog, E.M., Kõivomägi, M., Lucena, R., Mai, M.V., Hirschi, A., Kellogg, D.R., Loog, M., and Rubin, S.M. (2013). Cks confers specificity to phosphorylation-dependent CDK signaling pathways. Nat Struct Mol Biol 20, 1407–1414.

Meijer, L., Arion, D., Golsteyn, R., Pines, J., Brizuela, L., Hunt, T., and Beach, D. (1989). Cyclin is a component of the sea urchin egg M-phase specific histone H1 kinase. Embo J 8, 2275–2282.

Minshull, J., Blow, J.J., and Hunt, T. (1989). Translation of cyclin mRNA is necessary for extracts of activated Xenopus eggs to enter mitosis. Cell 56, 947–956.

Mochida, S., Maslen, S.L., Skehel, M., and Hunt, T. (2010). Greatwall Phosphorylates an Inhibitor of Protein Phosphatase 2A That Is Essential for Mitosis. Science 330, 1670–1673.

Morin, V., Prieto, S., Melines, S., Hem, S., Rossignol, M., Lorca, T., Espeut, J., Morin, N., and Abrieu, A. (2012). CDK-Dependent Potentiation of MPS1 Kinase Activity Is Essential to the Mitotic Checkpoint. Curr Biol 22, 289–295.

Moyle, M.W., Kim, T., Hattersley, N., Espeut, J., Cheerambathur, D.K., Oegema, K., and Desai, A. (2014). A Bub1–Mad1 interaction targets the Mad1–Mad2 complex to unattached kinetochores to initiate the spindle checkpoint. J Cell Biology 204, 647–657.

Musacchio, A., and Salmon, E.D. (2007). The spindle-assembly checkpoint in space and time. Nat Rev Mol Cell Bio 8, 379–393.

Nigg, E.A. (1995). Cyclin-dependent protein kinases: Key regulators of the eukaryotic cell cycle. Bioessays 17, 471–480.

Passmore, L., Booth, C., Venien-Bryan, C., Ludtke, S., Fioretto, C., Johnson, L., Chiu, W., and Barford, D. (2005). Structural analysis of the anaphase-promoting complex reveals multiple active sites and insights into polyubiquitylation. Molecular Cell 20, 855 866.

Peter, M., Nakagawa, J., Dorée, M., Labbé, J.C., and Nigg, E.A. (1990). In vitro disassembly of the nuclear lamina and M phase-specific phosphorylation of lamins by cdc2 kinase. Cell 61, 591–602.

Pines, J., and Hagan, I. (2011). The Renaissance or the cuckoo clock. Philosophical Transactions Royal Soc B Biological Sci 366, 3625–3634.

Pines, J., and Hunter, T. (1991). Human cyclins A and B1 are differentially located in the cell and undergo cell cycle-dependent nuclear transport. J Cell Biology 115, 1–17.

Qian, J., Beullens, M., Huang, J., Munter, S., Lesage, B., and Bollen, M. (2015). Cdk1 orders mitotic events through coordination of a chromosome-associated phosphatase switch. Nat Commun 6, 10215.

Qian, J., García-Gimeno, M., Beullens, M., Manzione, M., der Hoeven, G., Igual, J., Heredia, M., Sanz, P., Gelens, L., and Bollen, M. (2017). An Attachment-Independent Biochemical Timer of the Spindle Assembly Checkpoint. Mol Cell 68, 715–730.

Qiao, R., Weissmann, F., Yamaguchi, M., Brown, N.G., VanderLinden, R., Imre, R., Jarvis, M.A., Brunner, M.R., Davidson, I.F., Litos, G., et al. (2016). Mechanism of APC/CCDC20 activation by mitotic phosphorylation. Proc National Acad Sci 113, 2570–2578.

Reimann, J., Freed, E., Hsu, J.Y., Kramer, E.R., Peters, J.-M., and Jackson, P.K. (2001). Emi1 Is a Mitotic Regulator that Interacts with Cdc20 and Inhibits the Anaphase Promoting Complex. Cell 105, 645–655.

Rodriguez-Bravo, V., Maciejowski, J., Corona, J., Buch, H., Collin, P., Kanemaki, M.T., Shah, J.V., and Jallepalli, P.V. (2014). Nuclear Pores Protect Genome Integrity by Assembling a Premitotic and Mad1-Dependent Anaphase Inhibitor. Cell 156, 1017–1031.

Rowley, A., Choudhary, J.S., Marzioch, M., Ward, M.A., Weir, M., Solari, R., and Blackstock, W.P. (2000). Applications of Protein Mass Spectrometry in Cell Biology. Methods 20, 383–397.

van der Walt, S., Schönberger, J.L., Nunez-Iglesias, J., Boulogne, F., Warner, J.D., Yager, N., Gouillart, E., Yu, T., and contributors, scikit-image (2014). scikit-image: image processing in Python. Peerj 2, e453.

von Schubert, C., Cubizolles, F., Bracher, J.M., Sliedrecht, T., Kops, G.J., and Nigg, E.A. (2015). Plk1 and Mps1 Cooperatively Regulate the Spindle Assembly Checkpoint in Human Cells. Cell Reports 12, 66–78.

Schulman, B.A., Lindstrom, D.L., and Harlow, E. (1998). Substrate recruitment to cyclin-dependent kinase 2 by a multipurpose docking site on cyclin A. Proc National Acad Sci 95, 10453–10458.

Sørensen, C., Lukas, C., Kramer, E.R., Peters, J.-M., Bartek, J., and Lukas, J. (2001). A Conserved Cyclin-Binding Domain Determines Functional Interplay between Anaphase-Promoting Complex–Cdh1 and Cyclin A-Cdk2 during Cell Cycle Progression. Mol Cell Biol 21, 3692–3703.

Strauss, B., Harrison, A., Coelho, P., Yata, K., Zernicka-Goetz, M., and Pines, J. (2018). Cyclin B1 is essential for mitosis in mouse embryos, and its nuclear export sets the time for mitosis. J Cell Biol 217, 179–193.

Sudakin, V., Chan, G., and Yen, T.J. (2001). Checkpoint inhibition of the APC/C in HeLa cells is mediated by a complex of BUBR1, BUB3, CDC20, and MAD2. J Cell Biology 154, 925–936.

Sze, K., Ching, Y. P., Jin, D. Y., and Ng, I. (2008). Role of a Novel Splice Variant of Mitotic Arrest Deficient 1 (MAD1), MAD1β, in Mitotic Checkpoint Control in Liver Cancer. Cancer Res 68, 9194–9201.

Tatsumoto, T., Xie, X., Blumenthal, R., Okamoto, I., and Miki, T. (1999). Human Ect2 Is an Exchange Factor for Rho Gtpases, Phosphorylated in G2/M Phases, and Involved in Cytokinesis. J Cell Biology 147, 921–928.

Topacio, B.R., Zatulovskiy, E., Cristea, S., Xie, S., Tambo, C.S., Rubin, S.M., Sage, J., Kõivomägi, M., and Skotheim, J.M. (2019). Cyclin D-Cdk4,6 Drives Cell-Cycle Progression via the Retinoblastoma Protein’s C-Terminal Helix. Mol Cell 74, 758–770.

Vázquez-Novelle, M., Sansregret, L., Dick, A.E., Smith, C.A., McAinsh, A.D., Gerlich, D.W., and Petronczki, M. (2014). Cdk1 Inactivation Terminates Mitotic Checkpoint Surveillance and Stabilizes Kinetochore Attachments in Anaphase. Curr Biol 24, 638–645.

Visintin, R., Craig, K., Hwang, E.S., Prinz, S., Tyers, M., and Amon, A. (1998). The Phosphatase Cdc14 Triggers Mitotic Exit by Reversal of Cdk-Dependent Phosphorylation. Mol Cell 2, 709–718.

von Schubert, C., Cubizolles, F., Bracher, J.M., Sliedrecht, T., Kops, G., and Nigg, E.A. (2015). Plk1 and Mps1 Cooperatively Regulate the Spindle Assembly Checkpoint in Human Cells. Cell Reports 12, 66–78.

Welburn, J., Vleugel, M., Liu, D., Yates, J.R., Lampson, M.A., Fukagawa, T., and Cheeseman, I.M. (2010). Aurora B Phosphorylates Spatially Distinct Targets to Differentially Regulate the Kinetochore-Microtubule Interface. Mol Cell 38, 383–392.

Wieser, S., and Pines, J. (2015). The Biochemistry of Mitosis. Csh Perspect Biol 7, a015776.

Zachariae, W., Schwab, M., Nasmyth, K., and Seufert, W. (1998). Control of Cyclin Ubiquitination by CDK-Regulated Binding of Hct1 to the Anaphase Promoting Complex. Science 282, 1721–1724.

Zhang, S., Chang, L., Alfieri, C., Zhang, Z., Yang, J., Maslen, S., Skehel, M., and Barford, D. (2016). Molecular mechanism of APC/C activation by mitotic phosphorylation. Nature 533, 260–264.

Zuccolo, M., Alves, A., Galy, V., Bolhy, S., Formstecher, E., Racine, V., Sibarita, J., Fukagawa, T., Shiekhattar, R., Yen, T., et al. (2007). The human Nup107–160 nuclear pore subcomplex contributes to proper kinetochore functions. Embo J 26, 1853–1864.

